# Regulatory network hubs guide dynamic human lineage specification

**DOI:** 10.64898/2025.12.15.694070

**Authors:** Chikara Takeuchi, Sushama Sivakumar, Anjana Sundarrajan, Yihan Wang, Sean C Goetsch, Huan Zhao, Lei Wang, Mpathi Nzima, Minnie Deng, Kartik N Kulkarni, Lin Xu, Jun Wu, Bruce A Posner, Maria H Chahrour, W. Lee Kraus, Nikhil V Munshi, Gary C Hon

## Abstract

Transcription factors (TFs), including DNA binding proteins and epigenetic co-regulators, control the timing of gene programs during lineage commitment^1^. However, systematically defining the dynamic activities of the human genome’s nearly 2000 TFs during development remains challenging^2,3^. Here, we use Perturb-Seq to map the transcriptional impact of nearly all human TFs and a subset of enhancers during cardiomyocyte differentiation. Our results suggest that developmental regulatory networks are distributed across highly connected hub TFs^4–6^ that are dynamic across lineage and time, rather than organizing into top-down hierarchies directed by “master regulators”. We also identify distinct TF ensembles that regulate sequential decision points during cardiac cell-fate specification by coordinating lineage-specific activation with alternate lineage repression. Separately, analysis of TF-TF cooperativity uncovers a dynamic interaction between MEF2 family TFs^7^ and members of the Polycomb Repressive complex 1^8–10^ to execute alternative fate repression. Building on these regulatory interactions, we construct a deep learning transformer model to accurately predict perturbed TFs driving altered regulatory networks in patient-derived transcriptomes. Together, our results define the regulatory network architecture of human lineage specification and provide a platform for predicting TF function and interpreting disease mechanisms.

## Introduction

During development, cell-fate transitions are governed by biological networks including gene regulatory networks (**GRN**s)^4,5^. The architectures of these networks have important implications on their biological properties^11^. A central question in development is whether **GRN**s adopt a hierarchical architecture driven by cascades of master regulators or a more distributed architecture providing more robustness to perturbation^12,13^. However, much of the experimental evidence to support network topologies is derived from E. coli^14^ and yeast^15,16^, and evidence for determining network topology during the dynamic process of mammalian development remains limited^17^. Although human developmental GRNs and network topology can be inferred from genome-wide mapping studies^6^, the lack of interventions^2,3^ and temporal resolution limit current understanding.

Directed differentiation of human pluripotent stem cells (**hPSCs**) is a tractable model system for studying the mechanistic underpinnings of human development^18^. Transcriptional, epigenetic, and chromatin accessibility profiles exist across the time course of differentiation^19–25^ and these datasets have established key transition points and highlighted how epigenetic signatures are coordinated during lineage commitment. Importantly, human loss-of-function mutations in transcription factors (**TFs**), including DNA-binding proteins and epigenetic co-regulators, are associated with developmental disorders such as congenital heart disease (**CHD**)^26–34^. But, how perturbations of TFs alter gene regulatory networks during cardiac development or disease is not well investigated. Crucially, a more general question in development remains: How is lineage-specific activation coordinated with alternate-lineage repression during differentiation?

Addressing these questions requires a causal analysis of developmental GRNs. While computational strategies have been developed to infer GRNs, this remains a challenging problem^2,3^. An alternative experimental approach is to directly perturb TFs and measure their molecular outputs as gene expression changes. Stem cell differentiation is a particularly well-suited developmental model, since similar studies in animal models often result in embryonic lethality. We have previously optimized high-throughput Perturb-Seq^35–37^ in differentiating hPSCs^38,39^, which couples CRISPRi genetic perturbation with single-cell RNA-Seq to enable scalable interventional analysis with high content phenotypic readouts in dynamic models of human development.

Here, we perform CRISPRi Perturb-Seq to systematically knockdown 1983 TFs and 1267 enhancers during directed human cardiomyocyte (**CM**) differentiation, as one generalizable model. This interventional atlas identifies regulators of cardiac differentiation and reveals the dynamic architecture of TF-gene regulatory networks. We provide evidence that upstream control of cardiac differentiation is distributed across many transcriptional regulators, whereas downstream developmental programs are coordinated through a smaller set of stage-specific hub TFs such as MYOCD. By linking TFs to lineage programs with temporal resolution, we identify a series of sequential branching lineage decisions across cardiomyocyte differentiation that are regulated by specific sets of TFs. Our analysis also identifies combinatorial TF-TF interactions, some of which represent novel protein-protein interactions that function in lineage specification. Finally, to demonstrate how systematic perturbation atlases can aid in the interpretation of disease mechanisms, we use perturbation maps to develop Percoder, a machine learning model that predicts transcriptional drivers from patient-derived cardiomyocyte disease models.

## Results

### Systematic perturbation prioritizes 209 TFs required for cardiomyocyte differentiation

We performed Perturb-Seq on 1983 TFs (1747 DNA binding proteins and 236 epigenetic regulators) during directed differentiation from WTC11 iPSCs to Day 12 cardiomyocytes (**Figure 1a, Extended Figure 1a, Supplementary Table 1**). After filtering, this high-quality dataset consists of 765,351 cells (ave 3588 transcriptome unique molecular identifiers (UMIs), ave 1004 sgRNA UMIs, ave 1.9 sgRNAs called per cell) (**Supplementary Table 2**). We confirmed on-target repression efficiency (**Extended Figure 1b**) and replicate reproducibility among libraries (**Extended Figure 1c, 2**). Our differentiation protocol yielded cell states across cardiomyocyte differentiation, from early CMs (FN1+) to late CMs (TNNT2+) (**Extended Figure 1d**). Validating our screening approach, we observed depleted cell counts corresponding to several established regulators of cardiac differentiation (e.g. MEF2, GATA4, NKX2-5, HAND2, ISL1, and TBX5) (**Extended Figure 1e,f, Supplementary Table 3**).

**Figure 1.**
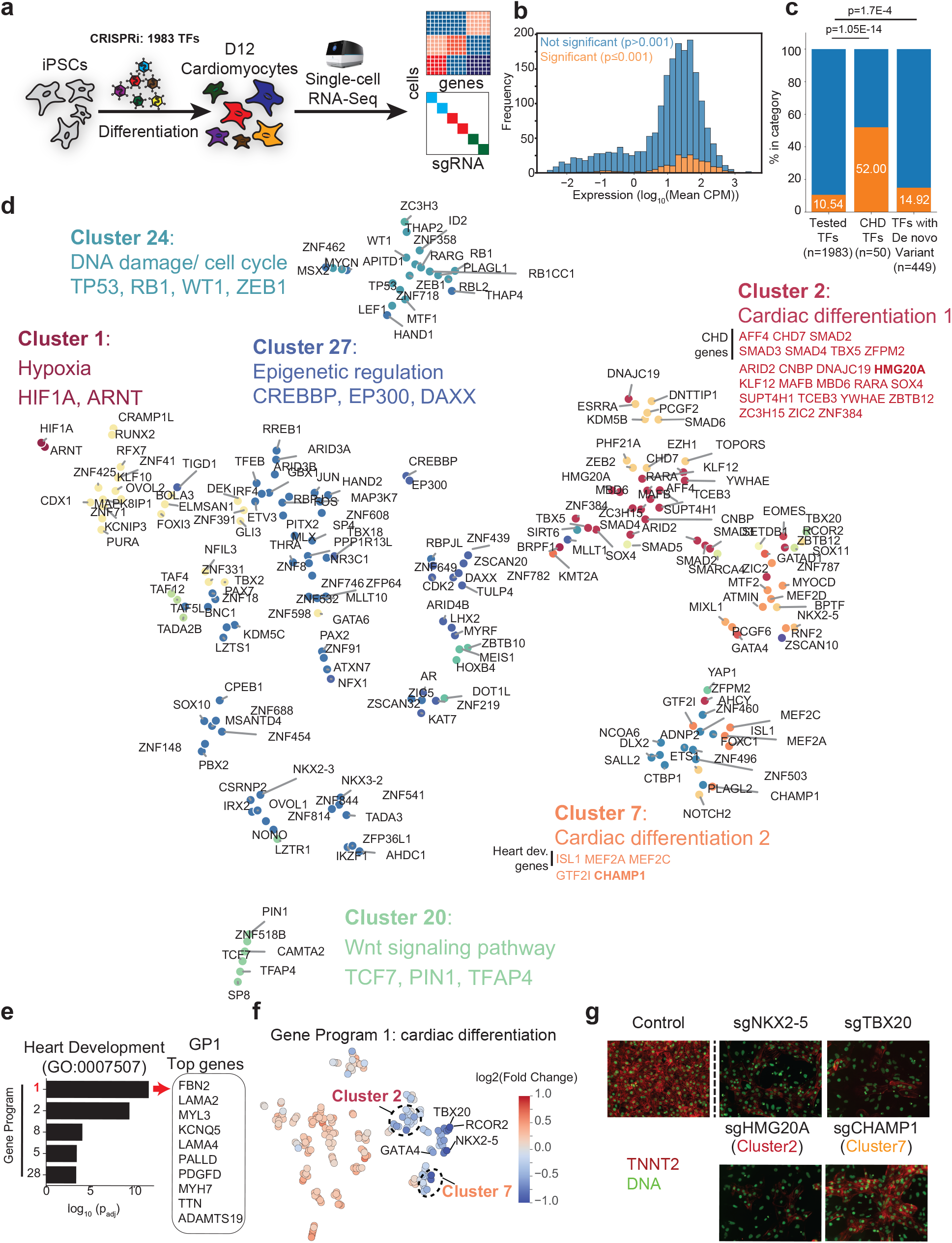
Perturb-Seq identifies 209 transcription factors regulating cardiac differentiation. a. Schematic of TF Perturb-Seq experiment design and subsequent analysis. b. Distribution of gene expression level for TFs perturbed in this study. Orange bars indicate significant TFs by energy distance. c. Enrichment of TFs for known congenital heart diseases genes (middle) and TFs with de novo variants (right). d. t-SNE embedding where each dot represents a significant TF perturbation(p=0.001). Colors indicate clusters. Labels of clusters and representative genes are annotated manually. e. Enrichment of heart developmental genes in each gene program. Top10 gene of Gene Program 1 is shown right. f. Feature plot showing GP1_card dev_ regulation for each perturbation. Embedding is the same as **Figure 1d**. g. Immunofluorescent staining of TNNT2 in iPSC-CM of Control KD, NKX2-5 KD, TBX20 KD, HMG20A KD and CHAMP1 KD.

We examined the Perturb-Seq dataset at multiple transcriptional resolutions. To identify TFs that impact global cell state, we adopted a principal components-based, energy distance (E-distance) metric reflecting transcriptome-wide phenotypes (**Extended Figure 3a**)^40^. We chose E-distance for initial triaging of TF hits given its ability to maintain the information content of single cell data from a complex differentiation regime in an unbiased and generalizable manner. Although E-distance is agnostic to biological mechanisms, it serves to prioritize TF candidates for downstream analysis. We identified 209 regulators that significantly impacted the transcriptome relative to non-targeting negative controls (**Extended Figure 3b-c,4, Supplementary Table 4-6**). Notably, even lowly expressed TFs could globally impact the transcriptome and hits were highly enriched in known CHD genes (p=1.05E-14) and in TFs with de novo variants (p=1.7E-4) (**Figure 1c-d, Supplementary Table 7**)^41–43^. Visually embedding similarities of TF transcriptome impacts by energy distance (**Extended Figure 3d**), we identify clusters encompassing the hypoxia pathway (Cluster 1: HIF1A, ARNT)^44^, DNA damage pathway (Cluster 24: TP53, RB1)^45,46^, transcriptional coactivators (Cluster 27: CREBBP, EP300)^47^, and the WNT pathway (Cluster 20: TCF7, PIN1)^48,49^ (**Figure 1d**). Interestingly, Cluster 2 contains regulators highly enriched in known CHD TFs, including TBX5^26,27^, SOX4^50^, CHD7^51^, and SMAD2-4^42,52,53^ (**Extended Figure 3e, Supplementary Table 8**), indicating shared transcriptional mechanisms. In addition, Cluster 7 represents regulators with known functions in heart development (**Extended Figure 3f**), including MEF2A^54^, MEF2C^55^, and ISL1^56^ as well as the poorly characterized transcription factors in cardiac biology such as GTF2I and CHAMP1.

To gain resolution, we applied consensus, non-negative matrix factorization (cNMF)^57^ to define gene programs (**GP**s) and examined GPs convergently regulated by multiple TFs (**Extended Figure 5a-b, Supplementary Table 9,10**). For example, we identified 24 TF perturbations that downregulate cardiac development GP1 (GP_card dev_), which is enriched with known heart development genes such as MYH7, TTN, MYL3 (**Figure 1e, Extended Figure 5c, Supplementary Table 11**). GP1_card dev_ is consistently down-regulated upon CRISPRi knockdown of TFs in Clusters 2 and 7 (**Figure 1f, Extended Figure 5d**), and immunostaining demonstrates reduced cardiomyocyte differentiation upon CRISPRi knockdown of HMG20A (Cluster 2), which has been associated with cardiac dysfunction in *Xenopus laevis*^58^, or CHAMP1 (Cluster 7), in which de novo mutations have been identified in patients with CHD (**Figure 1g, Extended Figure 5e-f**). Altogether, our systematic perturbation analysis identified a subset of 209 TFs that are required for CM differentiation, classified them by common downstream gene programs/transcriptional effects, and nominated additional uncharacterized TFs in cardiac development and disease.

### Hub TFs direct developmental gene programs in a distributed network

To address the question of whether the gene regulatory networks driving differentiation adopt a distributed or hierarchical topology^13,17^, we used Perturb-Seq data to construct GRNs based on perturbation-induced differential expression (**Figure 2a-b, Extended Data Fig. 6a**). This experimentally-derived GRN shares properties with other types of scale-free biological networks^16^ (**Figure 2b**; power-law parameter of 2.13), suggesting that distributed regulatory effects converge on a subset of highly connected nodes. As expected, genes with high in-degree (i.e., the number of TFs that regulate a given gene) are enriched for cardiomyocyte effector genes (e.g. TNNC1 and MYL3), many of which are co-regulated in GP1_card dev_ (**Figure 2b; right**).

**Figure 2.**
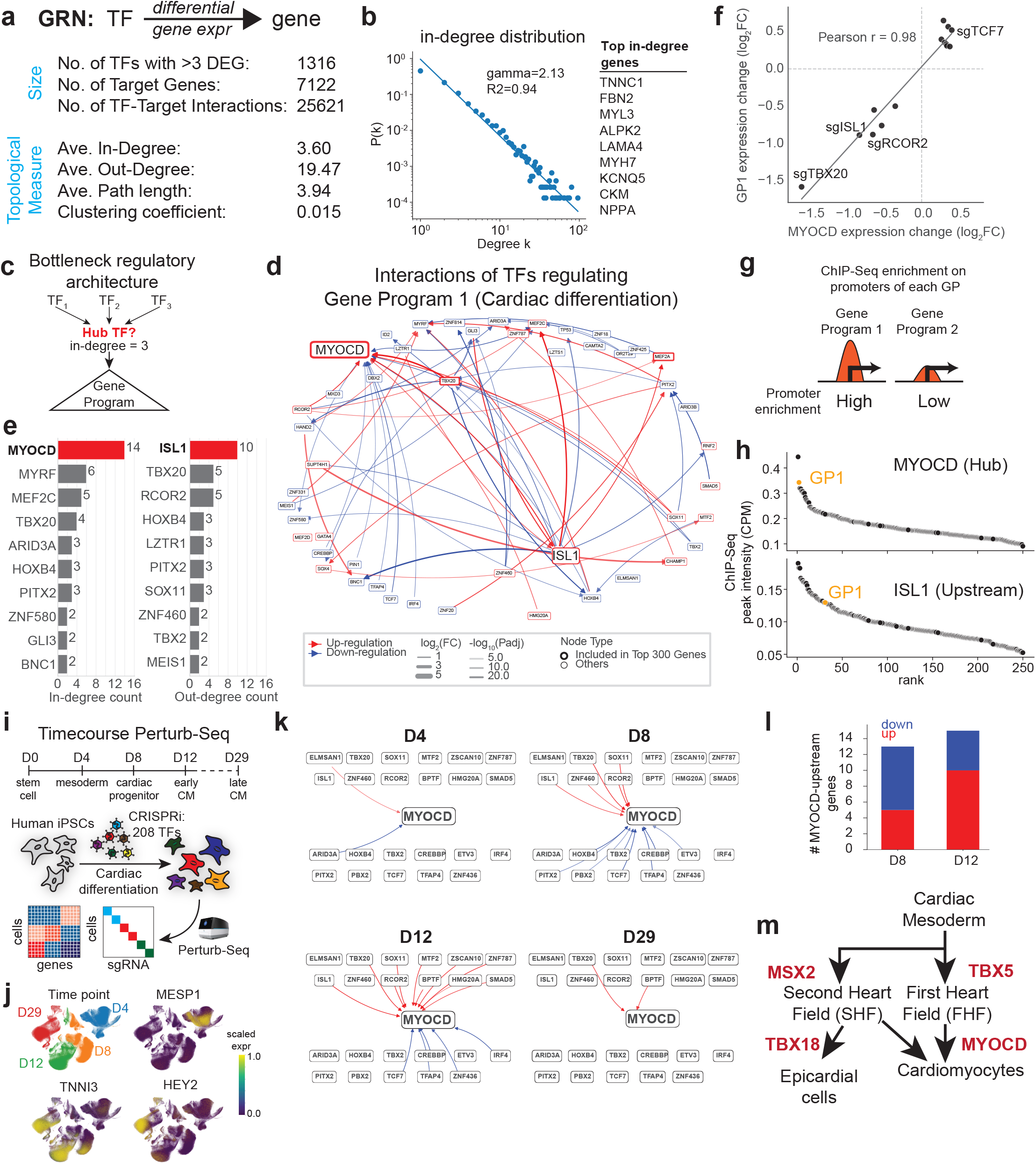
Hub TFs are regulatory interfaces for stage-specific developmental gene programs. a. Key statistics of the TF-gene regulatory network. b. Distribution of the in-degree distribution in the network. Top10 in-degree genes are shown at right. c. Schematic of strategy to identify a hub TF based on the in-degree in the TF-TF network. d. TF-TF regulatory network within GP1_card dev_ regulating TFs. Red arrows indicate positive regulation of the TFs and blue arrows indicate negative regulation of target TFs. e. Barplot showing in-degree count (left) and out-degree count (right) in TF-TF regulatory network. f. Scatterplot showing dose-response relationship of the MYOCD down-regulation and GP1_card dev_. g. Schematic of quantification method of TF enrichment on GP1_card dev_ promoters. h. Scatterplot showing enrichment of MYOCD and ISL1 ChIP-Seq signal on GP1_card dev_ promoters (TSS +/− 1kb). The x-axis indicates rank in all GPs, and the y-axis indicates mean CPM on promoters. i. Schematic of Time-course TF Perturb-Seq experiment design. j. Feature plot showing expression of the marker genes on UMAP. k. Dynamics of MYOCD regulators across cardiac differentiation. Only edges between upstream TFs and MYOCD are shown. l. Barplot showing in-degree edges of MYOCD and their regulation type. Red: up-regulation, Blue: down-regulation estimated by expression change pattern. m. Summary of the hub TF analysis in developmental-associated GPs.

We next explored the regulatory relationships of the TFs co-regulating GP1_card dev_. Systematic TF binding studies have proposed that TF-TF networks adopt a hierarchical structure with information bottlenecks^6^ (**Figure 2c**). To test this idea in the context of our developmental perturbation data, we counted TF-TF in-degree among gene program regulators (**Figure 2d**). Our analysis identified specific “hub TFs” that are regulated by many TFs and thus act as regulatory interfaces: among the 69 TFs associated with GP1_card dev_, 14 TFs regulate MYOCD (**Figure 2e; left**). Similarly, out-degree analysis confirms ISL1 as a key upstream regulator of cardiac differentiation^56^ (**Figure 2e; right**). Consistent with the notion that MYOCD is a regulatory interface used by other TFs to control cardiac differentiation, we observe a dose-dependent relationship between perturbation-mediated changes in MYOCD and expression of GP1_card dev_ genes (Pearson’s r = 0.98, n=14) (**Figure 2f**). We also observed significantly stronger occupancy of MYOCD at GP_card dev_ genes (rank 2)^59^ as compared to a non-hub TF, such as ISL1 (rank 31)^60^ (**Figure 2g-h, Extended Data Fig. 6b**). Together, these analyses support a distributed architecture in which upstream TFs influence differentiation programs through stage-specific hub TFs that interface with shared developmental gene programs.

### Hub TFs are dynamically regulated and stage-specific

Since prior models of human networks relied on data obtained from a fixed time point ^6,61^, we next wished to examine the dynamics of network topology throughout cardiomyocyte differentiation^22,23^ by performing time-course Perturb-Seq on the 209 TF hits across multiple cell states: mesoderm (Day 4), cardiac progenitor (Day 8), early cardiomyocyte (Day 12), and late cardiomyocyte (Day 29) (**Figure 2i-j, Extended Data Fig. 6c, Supplementary Table 12**).

For the MYOCD regulatory hub, we observed dramatic changes in network architecture across differentiation. While MYOCD is minimally regulated at the very early mesodermal (D4) and very late cardiomyocyte stages (D29), it is regulated by at least 13 TFs in the intermediate cardiac progenitor and early cardiomyocyte stages (D8-12) (**Figure 2k, Extended Data Fig. 6d**). Consistent with the role of MYOCD regulating cardiac commitment, the mode of MYOCD regulation also undergoes a switch from repression to activation between D8 (61.5% TFs repress MYOCD) and D12 (66.7% of TFs activate MYOCD) (**Figure 2l**).

Finally, to generalize these results, we extended this analysis to nominate hub TFs across several cardiac differentiation stages (**Extended Data Fig. 7a**). This analysis nominates hub TFs in cardiac mesoderm (Hand1), first-heart-field (Tbx5), second heart field (Msx2), and epicardial (Tbx18) gene programs (**Extended Data Fig. 7b-c**). Overall, these results indicate that a small number of hub TFs dynamically regulate developmental gene programs to drive cardiac cell type specification (**Figure 2m**).

### A series of branching lineage decisions drives cardiomyocyte differentiation

Cellular differentiation requires coordinated activation of specific lineage programs and repression of alternative cell fates^12,62^. To evaluate how TFs regulate lineage decisions during early human development, we aligned *in vitro* Perturb-Seq gene programs with *in vivo* spatial transcriptomic atlases of human embryos^63–65^. This approach enables us to define developmental gene programs in directed cardiac differentiation models, and to nominate their regulators (**Figure 3a, Extended Figure 8a-c**). For example, knockdown of GTF2I, ZBTB12, or RCOR2 decreased the activity of mesendoderm gene programs (GP45_ME1_, GP47_ME2_) while increasing an extra-embryonic mesoderm gene program (GP58_EXM_) (**Extended Figure 8d-g**). These results suggest that GTF2I, ZBTB12, and RCOR2 activate the mesendoderm lineage and repress the alternative extra-embryonic mesoderm lineage ^66^, an early lineage specification step required for proper cardiomyocyte differentiation.

**Figure 3.**
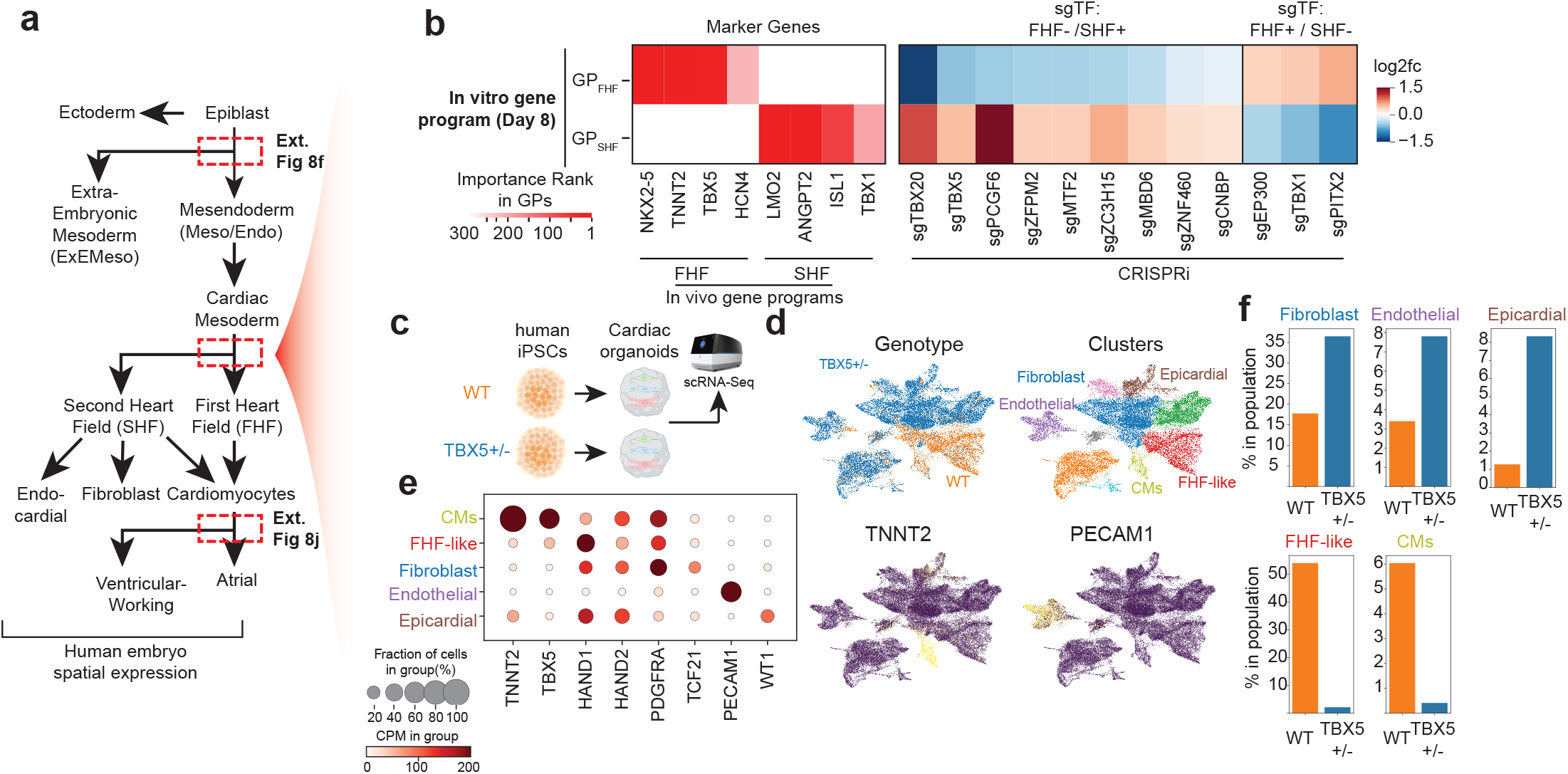
Time-course Perturb-Seq identifies transcription factors regulating lineage balance. a. Schematic of in vivo differentiation lineage from epiblast to distinct cell types. b. (Left) Heatmap showing key genes in GP10_FHF_ and GP32_SHF_ from D8 Time-course Perturb-Seq data. (Right) Heatmap showing changes of the GP usage at D8 cardiac differentiation. c. Schematic of experimental design for cardiac organoids. d. Feature plot showing genotype (upper left), leiden cluster and annotations (upper right), and cardiac marker (TNNT2, lower left) and endothelial marker (PECAM1, lower right). e. Bubble plot showing expression of the key marker genes in each cluster. Color indicates mean CPM expression, and size indicates percentage of expression in each cluster. f. Barplot showing quantification of the cluster distribution in each genotype.

Later during embryonic development, the first and second heart fields (**FHF** and **SHF**) emerge from cardiac mesoderm^67,68^ (**Figure 3a**). While the FHF primarily gives rise to CMs of the left ventricle, the SHF gives rise to endocardial cells (ECs), smooth muscle cells, fibroblasts, CMs of the right ventricle, atria, and outflow tract ^21,67,69^. We can detect the downstream consequences of this lineage branching by examining the status of corresponding GPs. By integrating our Day 8 Perturb-Seq experiment with spatial datasets from Carnegie stage 8-9 human embryos^63,64^, we identified GP10_FHF_ and GP32_SHF_ as FHF-like (TBX5+, HCN4+) and SHF-like (ISL1+, TBX1+) gene programs, respectively (**Extended Data Fig. 7a**). Knockdown of 20 TFs, including TBX5^70^ and several zinc finger TFs, significantly decreased GP10_FHF_ and reciprocally increased GP32_SHF_ (**Figure 3b, Extended Data Figure 8h, Supplementary Table 13-17**). Conversely, we identified 12 TFs, including Pitx2 and Tbx1, whose knockdown increased GP10_FHF_ and reciprocally decreased GP32_SHF_. To further test the idea that TBX5 activates FHF gene programs and represses SHF gene programs, we generated cardioids from wild-type and TBX5+/− hPSCs followed by single-cell RNA-Seq^39,71–73^ (**Figure 3c-e**). Cardioids with a 50% reduction in TBX5 exhibit a dramatic switch away from FHF cell states and a redirection towards cell states derived from SHF progenitors, such as cardiac fibroblasts^74^ (**Figure 3f**). Together, these results suggest that specific TF ensembles orchestrate FHF versus SHF specification by coordinating lineage-specific activation with alternative lineage repression.

Finally, during later CM subtype specification, we identified another set of TFs that impact the decision between atrial and ventricular myocyte lineages. For example, MYOCD knockdown significantly activated an atrial gene program (GP24_Atr_) by Day 29 and repressed the ventricular program (GP22_Vent_) (**Extended Figure 8i-j**). Importantly, we used an orthogonal supervised approach to annotate TF CRISPRi phenotypes by established lineage markers, demonstrating significant overlap with GP-based analyses (**Extended Figure 9a-c**). Taken together, we provide several examples from our Perturb-Seq datasets demonstrating that CM differentiation comprises a series of branching lineage decisions, during which specific TFs coordinate cell type-specific gene activation with alternative lineage repression.

### Cooperativity analysis identifies dynamic TF-TF interactions during CM differentiation

One key aspect of lineage specification remains poorly understood: how do TFs simultaneously coordinate lineage-specific gene activation and alternative lineage repression? Three potential mechanisms have been previously established: 1) cross-repression of lineage-specific TFs^75–77^, co-binding with repressive TFs at specific regulatory elements^62^, and 3) context-dependent co-repressor/co-activator recruitment^78^. Interestingly, signal-dependent chromatin-modifying complex exchange occurs in multiple contexts that impact differentiation, including the nuclear hormone receptor, Notch, Wnt, Hedgehog, and cAMP/CREB pathways^79–81^.Since chromatin regulators lack intrinsic DNA-binding activity and depend on TF-mediated recruitment, identifying the specific TF-chromatin regulator interactions involved remains particularly challenging. Thus, we developed a method for detecting distinct TF perturbations that affect common gene programs more than would be expected by chance (**Figure 4a, Supplementary Table 18**). Validating this approach, we found that HIF1A and ARNT, heterodimer partners in the hypoxia response^82^, co-regulate 29 gene programs (**Figure 4b**). This analysis also identifies frequent co-regulation of the MEF2 family of TFs (MEF2A, MEF2C, and MEF2D)^7,82^ with members of the Polycomb Repressive Complex 1 (PRC1; PCGF6, PCGF2, and RNF2)^8,9^ (**Supplementary Table 18**). These TFs regulate the expression of multiple gene programs associated with CHD (**Supplementary Table 18**). Interestingly, we notice a partitioning of gene programs regulated by distinct PRC1 complexes, where PRC1.2 (containing PCGF2 and RNF2) generally regulates a distinct set of gene programs from PRC1.6 (PCGF6). In contrast, TF usage is more varied: MEF2D only co-regulates PRC1.2 programs, while MEF2A shows enriched associations with both PRC1 complexes. (**Figure 4c**).

**Figure 4.**
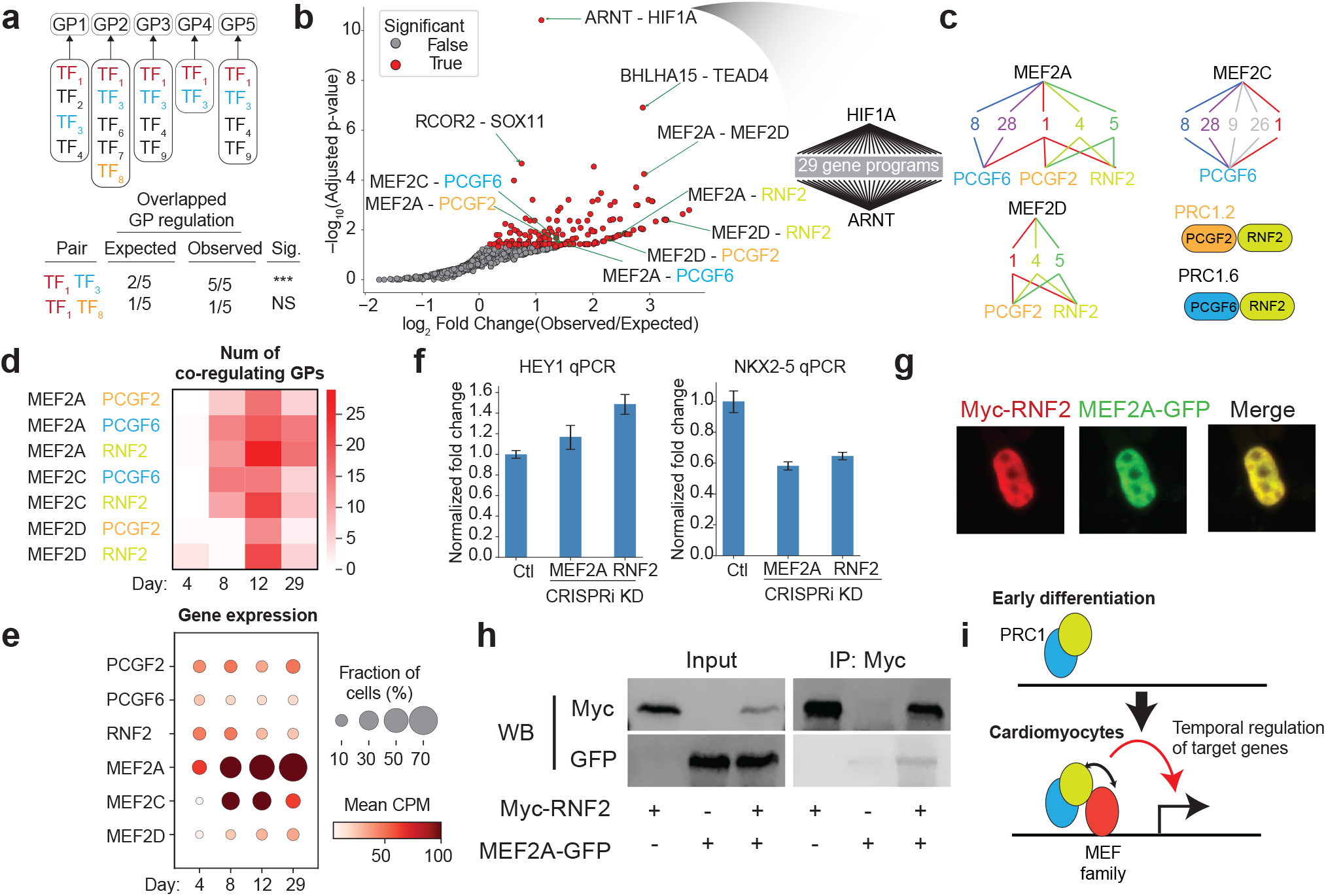
Co-regulation analysis identifies dynamic TF-TF interactions during differentiation. a. Schematic of co-regulation analysis of Gene Programs. We identified TF pairs that co-regulate multiple gene programs more often than expected by chance. b. Volcano plot showing occurrence of co-regulation between TF pairs. Significant pairs (p<0.05) are indicated as red. (right) Example of co-regulation of 29 gene programs between HIF1A and ARNT. c. An example of co-regulation between the MEF2 family and PRC1 complex genes. MEF2D is specifically associated with PRC1.2 (PCGF2), whereas MEF2A exhibits associations with both PRC1 complexes. d. Heatmap showing number of the co-regulating GPs between MEF TFs and PRC1 components at D4, D8, D12, D29. e. Bubble plot showing the expression of MEF2 TFs and PRC1 components in non-targeting gRNA cells in time-course Perturb-Seq library. The size of the dot indicates fraction of cells expressing marker genes, and color indicates mean CPM in non-targeting cells. f. qPCR results showing HEY1(left), and NKX2-5 (right) after MEF2A and RNF2 KD in ESC-CM. g. Immunofluorescence staining of Myc-RNF2 and fluorescence of MEF2A-GFP in the heterologous overexpression system. h. Co-immunoprecipitation of MYC-RNF2 and MEF2A-GFP, followed by Western blotting, to confirm physical interaction between RNF2 and MEF2A. i. Schematic of MEF2-PRC1 interactions revealed by co-regulation analysis.

Next, we examined MEF2-PRC1 interactions across our Perturb-Seq timecourse and observed that they are dynamic across differentiation, peaking in early cardiomyocytes (D12) (**Figure 4d**). This observation is consistent with a prior mouse study demonstrating that the PRC1.2 complex exchanges subunits in a stage-specific manner to promote cardiac differentiation by activating mesodermal fate specification and repressing alternative lineages^10^. These interactions vary with the expression of the MEF2 family of TFs, while the expression of PRC1 components is relatively constant (**Figure 4e**). To confirm that MEF2A and RNF2 (member of PRC1.2) coregulate shared gene programs, we performed qPCR in individual MEF2A or RNF2 CRISPRi CMs at day 12 to detect shared target genes. We confirm that HEY1 (marker of GP5) is modestly upregulated and NKX2-5 (marker of GP1_card dev_) is downregulated when MEF2A or RNF2 is repressed during CM differentiation (**Figure 4f**). Consistent with physical interaction, we observe that MEF2A and RNF2 protein expression overlaps in the nucleus (**Figure 4g**). To directly test whether RNF2 and MEF2A interact, we performed co-immunoprecipitation of Myc-RNF2 and MEF2A-GFP (**Figure 4h**).

Together, these results extend the utility of Perturb-Seq to identify complex gene-gene interactions in the context of development. Our findings support a model in which MEF2 family TFs dynamically cooperate with PRC1 complexes to coordinate cardiac developmental programs and repression of alternative-lineage-associated gene programs (**Figure 4i**) and provide a potential explanation for how Mef2 family TFs regulate distinct aspects of heart tube morphogenesis in mice^83^.

### An enhancer network controls TF expression

Enhancers controlling TF expression provide an additional layer to the cardiac lineage specification regulatory network ^39^ and harbor CHD-associated variants ^84^. Thus, we performed Perturb-Seq on a prioritized set of 1267 enhancers within TF regulatory neighborhoods (**Figure 5a, Extended Data Figure 10a-b, Supplementary Table 19,20**) and identified significant enhancers by energy distance analysis (**Extended Data Figure 10c, Supplementary Table 21-23)**. Enhancer perturbations for a given TF phenocopy promoter perturbations on a transcriptome-wide level (**Figure 5b, Extended Data Figure 11a-c, Supplementary Table 24**). To ensure comprehensive enhancer detection, we developed an approach to increase enhancer target gene sensitivity (**Extended Data Figure 11d-h, Supplementary Table 25**). A representative example from our data set is a CHD-associated variant in a MEIS1 enhancer (Enhancer781)^84^. Perturb-Seq of Enhancer781 and the MEIS1 promoter share modest transcriptome-wide similarity (**Figure 5c**), with upregulation of FBN2 and KDR and downregulation of TBX18 and GAP43, which we independently validate (**Figure 5d**). Importantly, the Enhancer781 variant also modulates the activity of a heterologous reporter gene^85^ (**Figure 5e**). Together, our data identify a layer of enhancers influencing the regulatory network for cardiac lineage specification and demonstrate that a CHD-associated MEIS1 enhancer alters its expression and downstream gene programs (**Figure 5f**).

**Figure 5.**
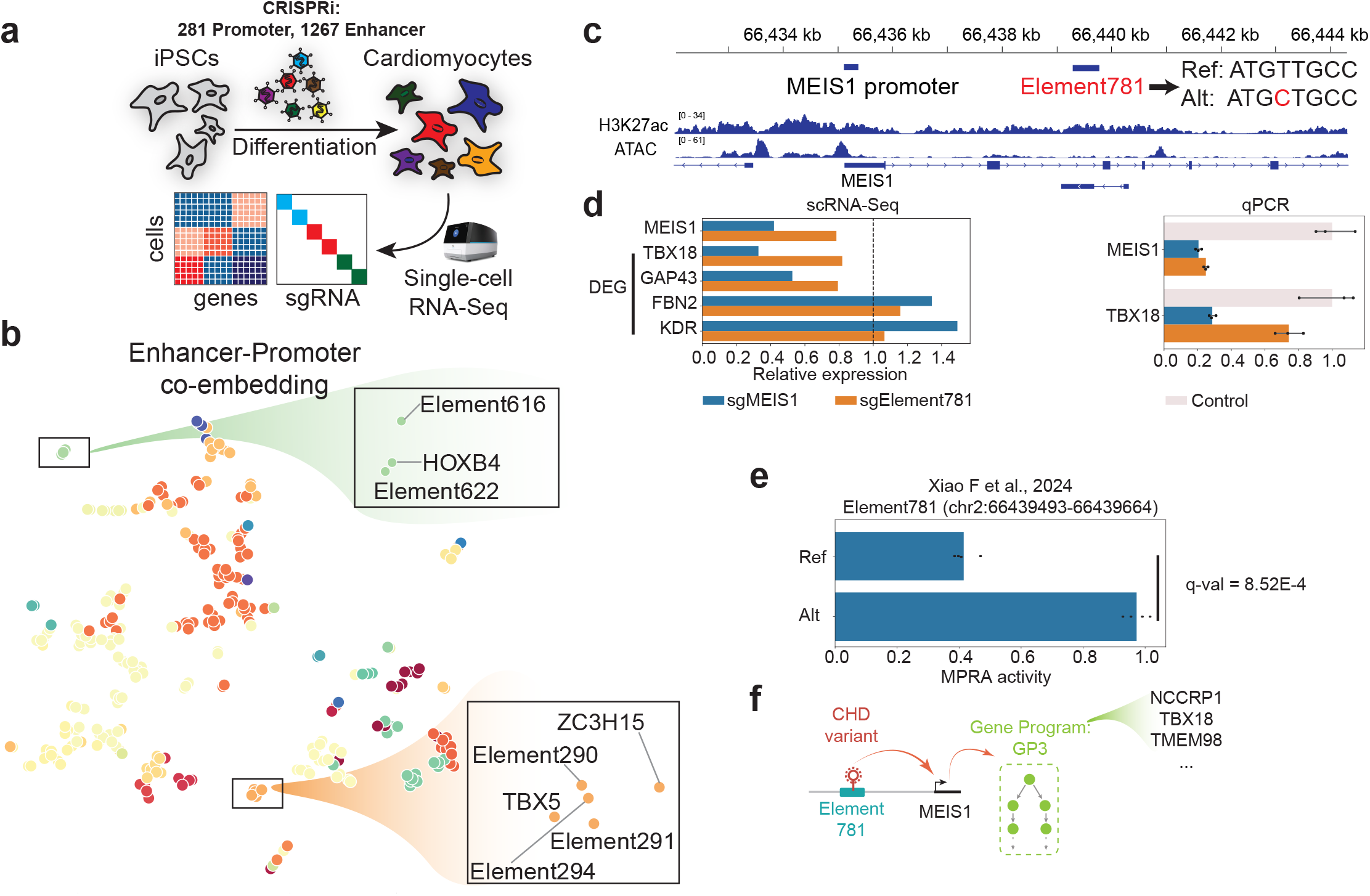
An enhancer network controls TF expression. a. Overview of enhancer Perturb-Seq experiment design. b. t-SNE embedding where each dot represents a promoter or enhancer perturbation. Colors indicate clusters. c. Genomic track showing the MEIS1 gene locus with enhancer Element781. Non-coding CHD de novo variants at Element 781^84^ are also shown. d. scRNA-Seq and qPCR results of the downstream impact by MEIS1 KD or MEIS1 enhancer (Element781) KD. e. MPRA result of reference and altered sequence at chr2:66439493-66439664 (overlapped with Element781). Y-axis indicates MPRA activity. Data From Xiao F et al., 2024^85^. f. Schematic summarizing a MEIS1 variant-enhancer-target-GeneProgram-Phenotype network.

### Percoder pinpoints disease-altered TF regulatory networks

Our interventional atlas of developmental networks provides a unique opportunity to infer candidate dysregulated regulators given transcriptional state and interpret disease mechanisms. As a proof-of-concept, we developed **Percoder** (a portmanteau of Perturb-Seq + Decoder) to predict underlying TF perturbations given unlabeled transcriptomes (**Figure 6a, Extended Figure 12a**). Percoder learns the transcriptional phenotypes of TF knockdown as a distribution over a set of cells using a Set transformer architecture^86^, which we use to predict perturbation labels (Methods) (**Extended Figure 12b**). Percoder predictions of 193 out of 210 perturbations (209 TF perturbations + non-targeting) achieve AUC > 0.75 (**Figure 6b**), with strong perturbations like sgNKX2-5 or sgGATA4 yielding near-perfect accuracy (**Figure 6c, Extended Figure 12c**). Instances of misclassification are biologically meaningful: ARNT and HIF1A (components of the hypoxia pathway), MEF2A and MEF2C (cardiac MEF2 family), EP300 and CREBBP (histone acetyl-transferases), and TBX5 and GATA4 (cardiac TFs and known interactors) (**Figure 6d**). Importantly, Percoder also performed well on out-of-sample data (**Extended Figure 12d-g, 13**)^87^ (Methods). Finally, we benchmarked the performance of Percoder against two pre-trained models, Geneformer (**Extended Data Figure 12h, Supplementary Table 26**) and SCENIC (**Extended Data Figure 12i-j, Supplementary Table 27**). Overall, our results indicate that Percoder effectively predicts TF perturbation responses for out-of-sample data.

**Figure 6.**
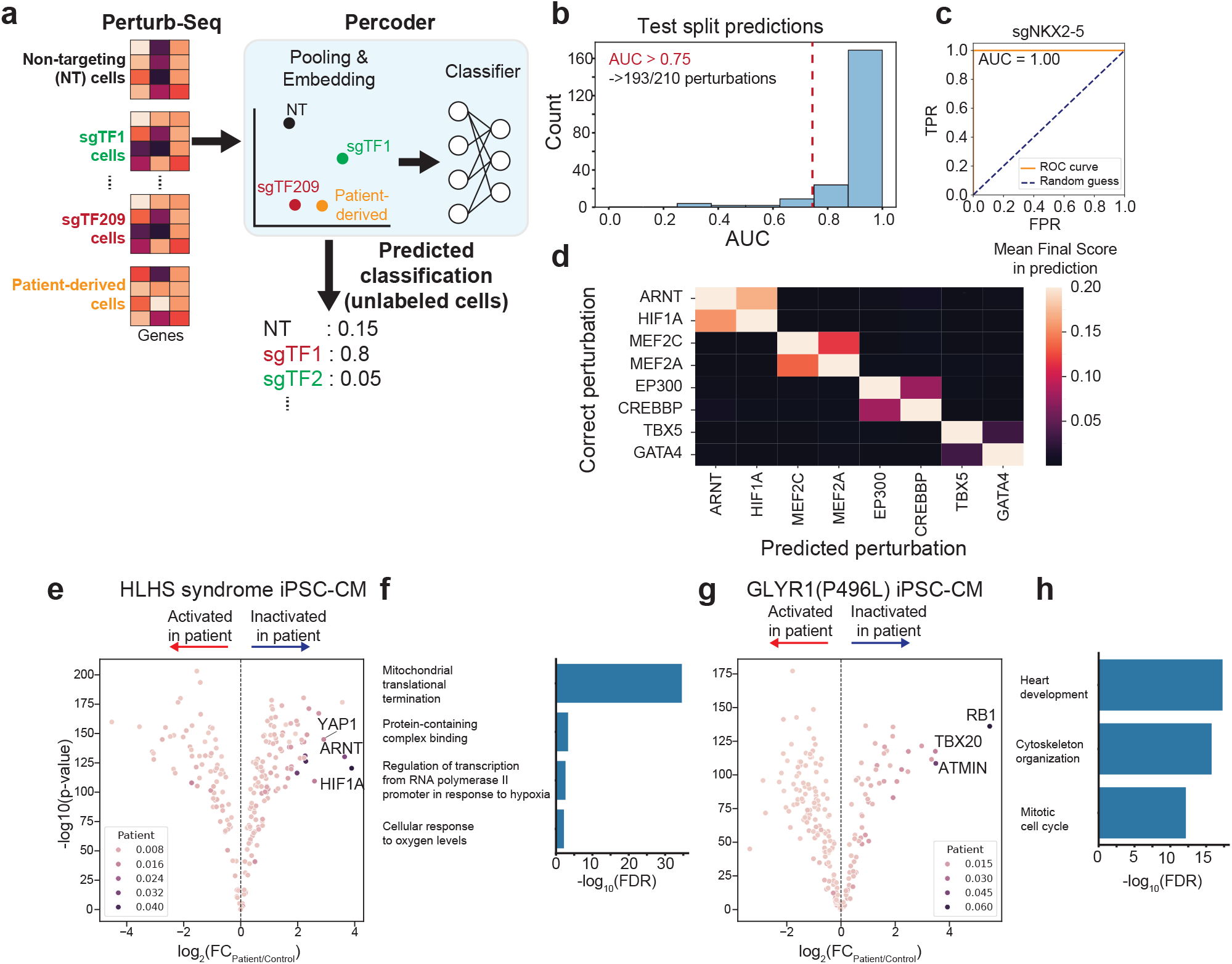
Perturbation modeling nominates TFs driving altered regulatory networks in patients. a. Schematic of Percoder framework, which uses a Set Transformer to pool and embed an input set of cells for the classification task of predicting perturbed genes. b. Histogram showing distribution of AUC per perturbation label. Red dashed bar indicates AUC=0.75 (threefold). c. Example ROC curve for the key cardiac TFs, NKX2-5. d. Heatmap of confusion matrix for TFs with related functions. e. We applied Percoder on scRNA-Seq data derived from HLHS patient-derived iPSCs differentiated to cardiomyocytes^88^. The volcano plot shows Percoder predictions of dysregulated regulators. Color indicates the Percoder score in the patient sample (not normalized relative to control). f. Gene Ontology analysis of differentially expressed genes between control and patient samples shown in PMID: 35395180. g. We applied Percoder on scRNA-Seq data derived from GLYR1 (P496L) patient-derived iPSCs differentiated to cardiomyocytes^60^. The volcano plot shows Percoder predictions of dysregulated regulators. Color indicates the Percoder score in the patient sample (not normalized relative to control). h. Gene Ontology analysis of differentially expressed genes between control and patient samples shown in PMID: 35182466.

Next, we used Percoder to predict dysregulated TFs from patient-derived iPSC-cardiomyocytes (iPSC-CMs) disease models. First, we analyze single-cell RNA-Seq datasets of iPSC-CMs derived from patients with Hypoplastic Left Heart Syndrome (HLHS)^88^. Percoder identified YAP1 as a top candidate dysregulated TF, consistent with the original published study identifying a defective YAP-regulated antioxidant response (**Figure 6e, Supplementary Table 28**). Importantly, the original study also observed significant dysregulation of genes in the hypoxia pathway in patients, which is consistent with Percoder’s prediction of ARNT and HIF1A as likely dysregulated TFs in patient transcriptomes (**Figure 6f**). Second, we apply Percoder to a single-cell RNA-Seq dataset of iPSC-CMs modeling a GLYR1 P496L point mutation found in CHD patients^60^ (**Figure 6g, Supplementary Table 29**). While the original study identified differentially expressed genes in heart development and mitotic cell cycle in the iPSC-CMs, Percoder goes further by predicting that these dysregulated networks are driven by TBX20, RB1, and ATMIN (**Figure 6h**). Third, analysis of scRNA-Seq data from iPSC-derived cardiomyocytes of a Hypoplastic Right Heart Syndrome patient^89^ identifies the myogenesis gene MTF1 as a potential driver (**Extended Data Fig. 12k, Supplementary Table 30**)^90^. Taken together, these results illustrate the utility of interventional atlases to understand mechanisms of diseases and identify potential targets for therapeutic intervention.

## Discussion

By comprehensively perturbing TFs in a model of human development, this study reveals several conceptual insights about the dynamic regulatory architecture of human stem cell differentiation. First, our GRN analysis supports the idea that upstream transcriptional regulators of developmental genes are distributed across many TFs, which promotes phenotypic redundancy during differentiation^4–6,16,61^, rather than being controlled by a hierarchical cascade of “master regulators”^91^. However, these distributed upstream regulators are also connected to “hub TFs”, such as MYOCD, that serve as integrators of regulatory signals. This distributed-but-convergent architecture may allow distributed upstream signals to drive developmental gene expression programs robustly. Hub TFs differ from classical master regulators (which sit at the top of a hierarchy) and directly regulate downstream differentiation genes. Interestingly, this is consistent with the idea that “middle managers” in a network regime are typically the executors of specific organizational programs^13,17^.

Second, by examining instances where multiple TFs frequently co-regulate shared gene programs, we identify TFs that function cooperatively in a developmental context. We identify interactions between the MEF2 family of TFs and members of the Polycomb Repressive Complex 1 that are dynamic throughout cardiac differentiation and may coordinate both activating and repressive effects on specific gene programs to direct cardiac lineage specification. The Polycomb Repressor Complexes are well-established to prevent alternative lineages at multiple points during differentiation^9,92^, and our analysis provides one example for how these complexes are dynamically recruited to execute their function. Interestingly, previous studies have linked MEF2 TFs with the PRC2 complex during cardiac development^93,94^ and fetal gene reactivation in heart failure^95^, but our results provide the first evidence that MEF2 TFs also function with PRC1 complexes.

This interventional atlas also facilitates our understanding of developmental diseases. While traditional genetic linkage studies have implicated several cardiac developmental TFs in familial CHD^26,31,96^, exome sequencing studies have recently identified de novo mutations for dozens of genes in sporadic CHD cases^41–43^. Many of the genes reaching statistical significance in these studies are also TFs and epigenetic regulators, including CHD7, KDM5B, KMT2A, PITX2, SMAD2, and SMAD6. Notably, our CRISPRi studies show that these genes are necessary for proper cardiac cell fate commitment and pinpoint specific defects. However, as CHD cohort sequencing studies are often underpowered, many potential CHD-relevant genes with de novo mutations have not reached statistical significance. In other words, these candidate genes are trending toward biologically meaningful associations with CHD, but larger cohorts will be necessary to justify these connections statistically. By attributing experimental evidence to the functions of TFs, our functional datasets provide additional support for several CHD candidate genes including: AHDC1, CHAMP1, CRAMP1L, EP300, KDM5B, KMT2A, LHX2, TADA2B, TBX18, and ZSCAN10. For example, exome sequencing identified a de novo nonsense mutation of CHAMP1 in a CHD proband (**Extended Figure 5f**)^41^, and our analysis shows that CHAMP1 knockdown significantly delays cardiomyocyte differentiation and phenocopies MEF2A and MEF2C knockdown (both of which are CHD TFs). Since patients with CHAMP1 mutations typically present with neurodevelopmental delay ^97^ and occasionally with heart defects ^98,99^, this example highlights how our resource can provide mechanistic data to support clinical observations and to reprioritize TFs with potential roles in CHD.

We demonstrate the application of experimentally defined regulatory networks with machine learning models to accurately predict perturbed TFs from the transcriptomes of unlabeled patient-derived cells. We anticipate that future maps of networks in other developmental contexts will expand our ability to predict the underlying TFs that are dysregulated in diseases across cellular contexts, which will promote virtual cell applications^100^. Reference maps of experimentally defined regulatory interactions like the ones presented in this study represent a foundational platform to understand the structure of regulatory networks in human developmental models, to model the functions of TFs, and to interpret the impact of genetic variants and cellular perturbations on development and disease.

Finally, we note some limitations of our study. First, our model system of human cardiac differentiation does not recapitulate all aspects of normal fetal development. For example, we use a two-dimensional, directed differentiation protocol that does not fully capture three-dimensional cardiac morphogenesis or the self-propagating nature of human embryogenesis. Second, the sensitivity of scRNA-seq is limited, which could lead to false negatives because only the most robust transcriptional effects can be detected at the expense of more subtle effects. Third, our network analysis was performed using perturbation phenotypes, rather than direct binding events, so it is possible that our analysis also incorporates indirect actions of individual TFs. Nevertheless, our approach balances the scale of Perturb-Seq with the ability to probe human development and generate high-quality datasets capable of accurate algorithmic training.

## Acknowledgements

We acknowledge the BioHPC computational infrastructure at UT Southwestern for providing HPC and storage resources that have contributed to the research results reported within this paper. This work is supported by NIH (UM1HG011996, R01HL136604, R01HL151650, R01HL176998, R35GM145235), the Welch Foundation (I-2103-20250403), and the Green Center for Reproductive Biology. We thank Eric Olson, Ning Liu, and Beverly Rothermel for critical feedback on the manuscript.

## Author contributions

**Chikara Takeuchi:** Methodology, Software, Formal analysis, Writing - Original Draft, Writing - Review & Editing, Visualization **Sushama Sivakumar:** Methodology, Validation, Investigation, Writing - Original Draft, Writing - Review & Editing, Visualization **Anjana Sundarrajan:** Methodology, Software, Formal analysis, Writing - Original Draft, Visualization **Yihan Wang:** Methodology, Formal analysis, Writing - Original Draft, Visualization **Sean C Goetsch:** Methodology, Investigation **Huan Zhao:** Investigation, Visualization **Lei Wang:** Investigation **Mpathi Nzima:** Data Curation **Minnie Deng:** Validation **Kartik N Kulkarni:** Validation **Lin Xu:** Funding acquisition **Jun Wu:** Funding acquisition **Bruce A Posner:** Funding acquisition **Maria H Chahrour:** Funding acquisition **W. Lee Kraus:** Writing - Review & Editing, Funding acquisition **Nikhil V Munshi:** Conceptualization, Writing - Original Draft, Writing - Review & Editing, Supervision, Project administration, Funding acquisition **Gary C Hon:** Conceptualization, Writing - Original Draft, Writing - Review & Editing, Supervision, Project administration, Funding acquisition

## Methods

### Experimental details

#### Cell culture

Human pluripotent stem cells (WTC11 and H9) are cultured as previously described^38^. In short, H9/WTC11 cells were maintained under feeder-free conditions in mTeSR plus medium (STEMCELL Technologies) and incubated at 37 □C with 5% CO2. For Perturb-Seq, we used the WTC11-CLYBL-dCas9-KRAB cell line^101^.

### Cardiac differentiation

WTC11 and H9 are differentiated to cardiomyocytes as previously described^38,39^. In short, 3 or 4 µM CHIR99021 supplemented CDM3 (RPMI 1640; 0.5 mg/ml human albumin, ScienCell OsrHSA; 211 µg/ml L-ascorbic acid 2-phosphate) for 48 hr followed by 2 µM Wnt-C59 supplemented CDM3 for 48 hr with subsequent media changes every 2 days with CDM3 alone. On day 12 post-differentiation, samples were washed with DPBS and dissociated to single cells with Accutase. For time-course experiments, WTC11 hiPSCs were differentiated into cardiomyocytes as described above. Cells were harvested at D4, D8, D12, and D29, with media changes performed every two days.

### Cardiac Organoid

Organoids were generated according to previous studies^72,73,102^. hiPSCs were seeded into 96-well plates at a density of 10,000 cells per well after achieving 70% confluency and dissociating with Accutase. Cells were resuspended in mTeSR Plus medium with ROCKi and incubated overnight. On day 0, media was changed to FlyAB(Ins), containing RPMI, B-27 with insulin supplement, FGF2, BMP4, LY294002, CHIR99021, and Activin A. Subsequent media changes were performed as follows: on day 1, with a refresh of FlyAB(Ins); after 36-40 hours, media was changed to BWIIFRa, which includes RPMI, B-27 with insulin supplement, BMP4, FGF2, IWP2, and retinoic acid. Media was refreshed every 24 hours for days 2-4 with BWIIFRa. On days 5/6, media was changed to BFI containing RPMI, B-27 with insulin supplement, BMP4, and FGF2. On day 7, the media was switched to maintenance media with RPMI and B-27, refreshed every 48 hours. Day 15 organoids were single cell dissociated using TrypLE, counted and used to construct scRNA-seq libraries.

### Design of sgRNA library

sgRNA sequences are from Replogle et al^103^. 6 gRNAs per region are used for each transcript. We included 600 non-targeting gRNAs and 600 targeting gRNAs as a control in the final library (**Supplementary Table 1)**. For enhancer Perturb-Seq, we designed gRNAs using FlashFry^104^. 10 gRNAs per target are used (**Supplementary Table 19**) based on the previous assessment of power-detection for enhancer Perturb-Seq ^105^.

### Perturb-Seq

Library construction, virus packaging, virus transduction and single cell RNA-Seq were performed as previously reported^38^. Briefly, DNA oligos (Twist Bioscience) are cloned into LentiGuide(10X)-BFP-Puro (LW203), and electroporated in Endura Duo competent cells (Lucigen). For virus packaging, 8µg of these plasmids are transfected in HEK293T with the 6µg of PMD2.G and 2µg of psPAX2 (Addgene ID: 12259 and 12260). The media is changed the next day and viruses are collected after 48hr of media change. These viruses are transfected to the WTC11-CLYBL-dCas9-KRAB cells at 80% confluency using the standard protocol, followed by BFP-positive sorting and 2 weeks culture. These cells proceed to the cardiac differentiation. For scRNA-Seq, cells were loaded onto a Next GEM Chip N with hash-tag oligo superloading^38,106^. Raw data were processed according to the manufacturer (10x Genomics, User Guide, CG000513).

### RNA extraction, cDNA synthesis and RT-qPCR

RNA extraction was done using Trizol according to previously published protocols^38^. cDNA was synthesized using lunascript RT (NEB) according to manufacturers instructions. qPCR was performed using SYBR-green master mix and specific primers to detect transcript levels. Biorad’s CFX 384 real time qPCR machine was used to quantify transcript levels. Used primer sequences are shown in a supplement data (**Supplementary Table 31**).

### Immunostaining, Immuno-precipitation and western blotting

For immunostaining, differentiated cardiomyocytes were grown on matrigel coated coverslips. Immunostaining was performed as previously described^107^. For immunoprecipitation and immunostaining in **Figure 4**, HeLa cells were transiently transfected with PCS2-Mef2A-GFP, PCS2-Myc-RNF2 constructs using Lipofectamine 3000 according to manufacturers instructions. MYC co-immunoprecipitation and immunoblotting was done as reported previously^108^. Input indicates 10% of lysates on western blot.

## Analysis details

### scRNA-seq data pre-processing, filtering low-quality cells and doublets

We used our in-house preprocessing pipeline Perturb-Seq-Processing-Pipeline available on GitHub (https://github.com/Hon-lab/Perturb-Seq-Processing-Pipeline/). In short, we mapped the transcriptome to hg38 with Cell Ranger (version 7.0.0, 10x Genomics). Cells with low UMI count or higher than 20% mitochondrial content were removed, as well as genes with fewer than 1 counts. Also, gRNAs and HTO are assigned to cells using FBA^109^. Singlet cells (only 1 HTO) with at least 1 gRNA are used in the following analysis. For visualization, we applied UMAP based on 50 principal components from principal component analysis (PCA).

### Energy distance overview

To compare the effect of perturbations and associations between perturbations, we use energy distance analysis^40,110,111^. To balance computational cost and precision, we applied principal component analysis(PCA) (described above), and kept the top 50 principal components for the downstream analysis.

For samples *x*_1_,*x*_2_,…,*x*_*n*_ from X and *y*_1_,*y*_2_,…,*y*_*n*_ from Y, energy distance E(X,Y) is calculated in the following formula:

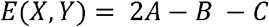

where A, B, and C are simply averages of pairwise euclidean distances of population (1) X and Y, (2) X and X and (3) Y and Y:

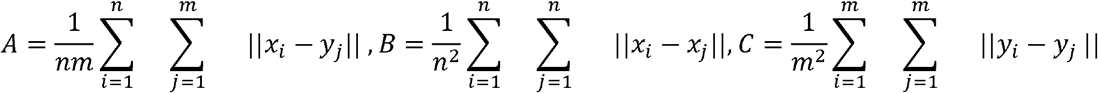

∣ ∣ · ∣ ∣ denotes the Euclidean norm of its argument. We implemented this calculation using pytorch to boost speed with the Tesla V100 32GB (see code availability).

### Removing outlier gRNAs

The first step of analysis is removing outlier gRNAs within each target. Although CRISPRi has less off-target activity, we observed some examples of off-target activity gRNAs and weak gRNA, that affect associations between perturbations (**Extended Data Fig. 3**). Some methods filter out cells based on target gene expression level^112^ or differential expressed genes^113^. However, these methods are not appropriate for our setting because (1) gene expression is highly changing over time in differentiating dataset, and (2) cell states are not uniform even with non-targeting gRNAs and (3) quantification of transcription factor expression is sometimes difficult especially for transient or lowly expressed but important genes. Therefore, we propose a new algorithm based on energy-distance. Assumptions of this algorithm are (1) cells with the same gRNA have the same off-target and on-target (repression efficiency) effects (2) cells with outlier gRNAs have a different distribution in the embedding space. These assumptions motivate us to filter gRNAs based on metrics which do not depend on target gene expression level. This algorithm consists of three steps.

1. *Filtering gRNA based on # of gRNAs and cells* If less than 20 cells are assigned to a gRNA, this gRNA is not considered in the following analysis because it would be noisy to use a small number of cells in the analysis. After this filtering based on cell number, if only 1 or 2 gRNAs are left after drop-off or filtering out gRNAs with a small number of cells, all gRNAs are kept. If not (if the number of gRNAs is more than 2), proceed to step 2.
2. *Test equal distributions among gRNAs using DISCO* First step is to distinguish whether outlier gRNAs are included in a given gRNA set. The purpose of this process is to judge outlier gRNAs are included in the gRNAs or not. Then, we tested the equal distribution of all gRNAs using DISCO (distance components), which is a nonparametric extension of ANOVA using energy distance. Let *A*_*i*_ ={*a*_*i*1_, *a*_*i*2_,…, *a*_*in*_,} is a set of cells with the gRNA i. To calculate F statistics of DISCO:

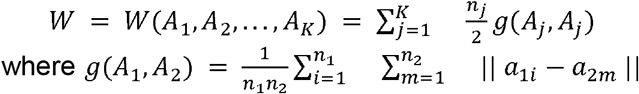

And total dispersion T is 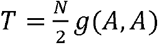 where A is the pooled sample of(*A*_1_,*A*_2_,…,*A*_*K*_). Then, the between-sample energy statistic S is calculated as*S*=*T−W*.

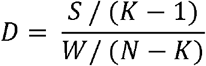

This *D* is an analogue of F statistics in ANOVA and calculated empirically using a permutation test. Practically, we shuffled the label of gRNA assignment within tested gRNAs (we use 1000 permutations for each target). p-value is calculated by the following formula.

*p =* #(*D*_*permute*_ > *D*_*observe*_) /#(*permutation*) Where *D*_*permute*_ represents for Dstatistics after permutation of gRNA labels and *D*_*observe*_ represents observed D statistics with right labels. If p-value of DISCO is more than 0.05, gRNAs per target is judged as all equally distributed. In other words, each gRNA behaves similarly. If p-value is less than 0.05, these gRNAs are likely to contain outliers within these, therefore proceed to the “Finding outlier gRNAs” step.

#### (3)Finding outlier gRNAs

The assumption here is that “clean” gRNAs, which show no off-target activity and strong repression of targeted regions, are distributed similarly. Thus, the energy distance between cells with distinct clean gRNAs will be smaller than between outlier gRNAs. This idea is similar to the Local outlier factor analysis of k-nearest neighborhood method, with some modification: (1) to calculate p-value in an unbiased way and (2) to detect outlier gRNA in small numbers of samples (max 6 gRNAs per target).

1. Let g_1_, g_2_, g_3_, g_4_ be gRNAs targeting a given gene, and X_m_ = {x_m,1_, x_m,2_, …, x_m,n_} are cells with g_m_ detected. Among these gRNAs, there may exist outlier gRNAs with distributions that significantly differ from other gRNAs. We propose the algorithm below to detect these outlier gRNAs.
2. Generate all possible pair combinations of gRNAs (targeting the same gene). For instance, given five gRNAs [A, B, C, D, E], we consider pairs such as [(A,B), (E)] or [(A,C,D), (B,E)] where both the total number and the combination of labels within each pair are arbitrary. We denote the first group as G1 and the second group as G2. For example, If we test the group of [(A,C,D), (B,E)], G1 is (A,C,D) and G2 is (B,E) (**Data S1**).
3. Calculate the Energy distance between G1 and G2 for all possible combinations.
4. Rank the calculated Energy distances in ascending order. If a gRNA is an outlier, the distances between groups containing this label and other groups should be larger, resulting in higher ranks for combinations including the outlier label (**Data S1**). If there is no outlier gRNA, combinations containing any given gRNA will be equally distributed or skewed to lower rank. Therefore, the frequency of combinations where a specific gRNA is skewed to higher rank can be used as a metric of how much this gRNA is outlier. We test this hypothesis using a hypergeometric test for each label, specifically examining the probability of combinations containing that label falling into the upper 50% of ranks. An adjusted p-value after Bonferroni correction is less than 0.05 is detected as outlier gRNAs and filtered out in the following analysis unless mentioned.

## Detecting significant hit regions

After removing outlier gRNAs we performed a permutation test using energy distance as described^40,110^. In short, we calculate energy distance between cells with non-targeting control gRNAs and cells with targeting gRNAs. For 1 batch, we randomly subsampled 2000 non-targeting cells to faster computation, and 1000 permutations were performed per batch. We repeated this step for 20 batches to avoid a bias effect on choice of background non-targeting cells. In total 20000 permutations were tested per target. To note, p-values are consistent among different sets of 2000 non-targeting cells (**Supplementary Table 5, Supplementary Table 23**). In this study, perturbations whose permutation-based p-value is less than 0.001 are referred to as significant hits.

## Association analysis, clustering, and embedding of perturbations

Within significant TF hits, we calculated energy distance between perturbations (**Extended Data Fig. 3d**). Clustering of these significant hits was performed in the following method. First, distance matrices from energy distance analysis are embedded in high dimension space with Isomap (implemented by sklearn.manifold.Isomap). This step is important because biologically associated TF pairs such as EP300 and CREBBP (both are histone acetyltransferase), HIF1A and ARNT (both are hypoxia pathway genes) significantly show smaller distances than other TFs, whereas larger distances between perturbations generally have little information, and Isomap is able to retain information of local structure in distance space. After embedding in high dimension space with Isomap, we clustered perturbations using affinity propagation (implemented by sklearn.cluster.AffinityPropagation) in Isomap embedded space to find clusters in an unbiased way without defining the number of clusters.

### Comparison between CHD genes and de novo variants

The list of congenital heart disease genes was obtained from CHDgene^114^. Among CHD genes, we extracted overlapped genes with our tested TFs, and calculated p-value of enrichment using a hypergeometric test. For de novo variants, we obtained the list of de novo variants from multiple studies^41–43^.

### Gene programs analysis

Gene program analysis was performed with consensus non-negative matrix factorization (cNMF)^57^. cNMF was run for 100 iterations for top 2000 highly variable genes using 0.1 as clustering metric. The optimal k (k=250) was identified based on unique gene ontology terms^115^, significant perturbations identified, and recovery of a rare perturbation (REST) as a positive control.

#### Significant perturbation calculation

cNMF assigns a score for each cell for every program. The mean score was calculated for each target for each program. Similarly the mean score was calculated for non-targeting control. Log_2_ fold change was calculated between targeting and non-targeting for every program. Mann-Whitney U test was performed to determine significance and Benjamin-Hochberg for adjusted p-value. A cutoff P-value of 0.001 was used based on the olfactory receptor genes as negative control^116^.

### ChIP-Seq analysis

ISL1 ChIP-Seq data from day6 WT hiPSC ^60^, and MYOCD ChIP-Seq data from day5 WT hiPSC^59^ are used in this study. We used custom scripts to calculate signal enrichment within ±1 kb of the promoters for the top 100 genes. For genes with multiple promoters, the MANE Select promoter was used.

### Annotation of the gene program

Cardiac differentiation gene programs (**Figure 1f**) and cardiomyocyte cell type (**Figure 2m**) are annotated by importance of the marker genes for each gene program based on the previous literature ^67^. For gene program annotation of Extraembryonic mesoderm/Mesendoderm and sub-type of cardiomyocytes (**Extended Data Fig. 7a,b**) analysis, we annotate gene programs by overlapping with marker genes in reference map^64,65^.

### SCENIC analysis

We used SCENIC^117^ algorithm to identify gene regulatory networks (regulons) activated in each fibroblast cluster with raw count matrix as input. Briefly, the co-expression network was calculated by GRNBoost2 and the regulons were identified by RcisTarget. Next, the regulon activity for each cell was scored by AUCell. Changes of usages of each regulon is calculated by fold changes of mean usage in the perturbed cells and non-targeting cells, and p-values are calculated by Mann-Whitney U test. Rank of the target regulon is calculated by in-house pipeline (See Code availability)

### Geneformer Analysis

The raw count matrix of the **Figure 1** dataset (209 TF perturbations) and **Figure 5** dataset (83 TF perturbations) are tokenized by the geneformer tokenize_data function. This tokenized data was embedded in the Geneformer-V2-316M model. To evaluate the data in a manner consistent with the in silico perturbation analysis described in the original study^118^—where rank-altered token embeddings were compared against other datasets via cosine similarity—we implemented the following procedure. First, reference embeddings were established by calculating the mean embedding of perturbed or non-targeting cells from the Figure 1 dataset. Similarly, the mean embedding was computed for cells within the Figure 5 dataset. Finally, we assessed the similarity between the Figure 5 embeddings and each Figure 1 reference using cosine similarity to identify the most closely aligned profiles.

### Identification of TF-TF interactions from gene program analysis

To estimate interaction effect sizes, we generated a degree-preserving bipartite null model using 10000 permutations. This method preserved both TF frequency across programs and the number of TFs per program while randomizing TF-program assignments. Interaction effect sizes were quantified as the log □ ratio of the observed co-occurrence count to the mean null co-occurrence count. Statistical significance of TF-TF overlap was assessed using a hypergeometric test, followed by Benjamini-Hochberg correction. Interactions with adjusted p-value < 0.05 were considered significant.

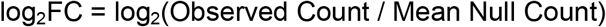

To ensure biologically meaningful interactions, we applied two additional filters: (1) Interactions with an observed count <2 were removed; (2) Interactions between two promoters of the same gene were excluded.

### Differential expression (DE) analysis

pySpade (version 0.1.6) is used to calculate differentially expressed genes^119^. pySpade ‘process’ function takes the output files from our in-house pre-processing pipelines. The randomization method of the DErand function is set to be ‘equal’ for all the datasets. Complementary cells are used in the background sets except time-course datasets. Non-target cells are the background in time-course datasets. All hits reported are FDR < 0.1, with fold change >20% as cutoff, and genes expressed in at least 5% of cells.

### Percoder

#### Architecture of Percoder

The architecture of the Percoder is based on the Set Transformer architecture. A set of N cells is denoted as 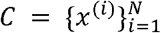 where *x* ^*(i)*^ ∈ *R*^*G*^ represents G-dimensional embedding of the i-th cell (e.g., raw or normalized counts, PCA components, etc.). First, each cell embedding *x* ^*(i)*^ is passed through a multilayer perception (MLP),*f* _*cell*_, with ReLU activation, followed by a Dropout layer. This transforms the input from a G-dimensional vector to a G’-dimensional vector, so *x* ^*’(i)*^ ∈ *R*^*G’*^.

Next, the transformed cell set 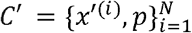 is processed by a **Set Attention Block (SAB)**. The structure of this SAB reflects the Set Transformer++ architecture, incorporating residual connections:

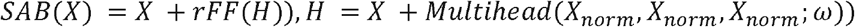

Where *X*_*norm*_=*Layern Norm*(*X*), and *ω* is the parameter for the Multihead attention layer. The output is denoted as *Z*=*SAB*(*C*^*’*^).

After the cell embeddings are processed by the attention mechanism, they are aggregated into a single vector using a Pooling by Multihead Attention (PMA) layer:

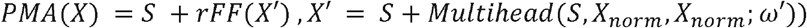

where *X*_*norm*_=*Layern Norm*(*X*)and S ∈*R*^*G’*^ is a learnable seed vector. *S* is seed vector with G’ dimension, and *ω*^*’*^ is the parameter for the Multihead attention layer.

The resulting output vector, *V* = *PMA*(*Z*)∈*R*^*G’*^ is then passed through a final MLP,*f* _*class*_, with ReLU activation, and a Softmax layer to predict the probability for each class (i.e., perturbation).

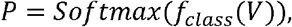

The final outputis *P* ∈ *R* ^210^ a probability distribution over the 210 classes.

#### Optimization of cell set-based classification problem

The reverse-perturbation problem is discussed in the previous foundation model studies^120,121^. The innovations in Percoder to predict perturbations is the use of sets of the cells for training, instead of single cells for training. We observe significant improvement of prediction accuracy if we use more cells for training/validation (**Extended Data Fig. 12b**). This indicates that heterogeneity in Perturb-Seq data is meaningful in a biological way to distinguish perturbations. A recently published perturbation prediction model called STATE also uses a similar approach in terms of using sets of the cells as input^122^. In the STATE model, the prediction accuracy from a set of cells dramatically outperforms one from using the mean of cells. This also suggests that heterogeneity of the cell transcriptome encodes biological information. To achieve best prediction accuracy, we noticed that Online Label Smoothing^123^ significantly improved final model prediction (**Fig. 6d**). Online label smoothing can learn conceptually similar labels. It is true that some of the TFs are known to form protein complexes (e.g. HIF1A and ARNT), or work in the same pathway (CREBBP and EP300). We speculate that the loss function of OLS helps to distinguish these similar concepts, and leads to better classification accuracy.

#### Optimization of generalization

To achieve the generalizability of Percoder, we embedded scRNA-Seq using the conditional variational auto encoder (cVAE) using scVI^87^. During hyperparameter tuning, we noticed that some of the parameters are critical to generalizable embeddings. First ZINB modeling of transcriptome outperforms NB modeling. Second, not highly-variable genes but using all of the genes improves the final prediction accuracy in the out-of-sample experiment. These results indicate that ZINB models can express single-cell transcriptomes well and technological/biological effects are expressed even in the lowly expressed genes.

## Data and code availability

The complete FASTQ data, sgRNA sequences, differential expression tables and all relevant metadata are available on the IGVF Consortium Data Portal (https://data.igvf.org/). The data used throughout this paper can be accessed through the following accession numbers and links: https://data.igvf.org/analysis-sets/IGVFDS6332VCTO/ https://data.igvf.org/analysis-sets/IGVFDS8939YRXO/ https://data.igvf.org/analysis-sets/IGVFDS1245HFTX/

scRNA-Seq data for cardiac organoid from hiPSC presented here are available for download at GEO under the accession number GSE330353.

Software and codes related to this study are on GitHub: https://github.com/Hon-lab/TFperturb_CM https://github.com/Chikara-Takeuchi/Percoder https://github.com/Hon-lab/Perturb-Seq-Processing-Pipeline

Any additional information required to reanalyze the data reported in this paper is available from the lead contact upon request.

## EXTENDED DATA FIGURES

**Extended Data Fig. 1**

a. Breakdown of the 1983 transcriptional regulators perturbed in this study.

b. Knockdown efficiency of target genes in Perturb-Seq. Lowly expressed genes (Log10(CPM_non-target_)<0.5) are excluded in the plot. The density plot is shown on the right.

c. Feature plot of the transcriptome UMIs per cell.

d. Feature plot of the late-stage CM marker TNNT2 and the early stage CM marker FN1.

e. Heatmap showing the Pearson correlation of gRNA abundance at Day 0, 7 (by bulk amplicon sequencing), 12 (by scRNA-Seq).

f. Comparison of sgRNA abundance: (left) Day 0 vs Day 7 (bulk vs bulk); (right) Day 0 vs Day 12 (bulk vs single-cell).

**Extended Data Fig. 2**

Feature plot illustrating the cell distribution for each of the 28 TF Perturb-Seq libraries. Library 5 recovered fewer cells because a lower volume was recovered during droplet generation.

**Extended Data Fig. 3**

a. Schematic of energy-distance analysis

b.Bar plot showing significant genes called at different p-value cut-offs. Orange bars indicate the number of hits called for Olfactory Receptor genes (OR genes, which represent targeting negative controls).

c. Bar plot showing enrichment of OR genes in the significant hits at different p-values.

d. Heat map matrix of the energy distance between 209 significant perturbations.

e. Enrichment of known CHD genes in each cluster. The names of TFs in Cluster 2 are listed.

f. Enrichment of heart developmental genes in each cluster. The names of TFs in Cluster 7 are listed. Orange TF names indicate heart developmental genes within the cluster.

**Extended Data Fig. 4**

a. An example of using energy distance analysis to filter out poorly performing sgRNAs. One sgRNA targeting HIF1A (top, 62162262.23-P1P2-1) does not correlate with other sgRNAs targeting HIF1A and is an outlier. One sgRNA targeting ARNT (150848808.23-P1P2-2) has weaker correlation with other sgRNAs targeting ARNT.

b. Schematic of our strategy to filter out poorly performing sgRNAs. (left) We are given sgRNAs (g1, g2, g3, g4) targeting a given gene. Each dot represents a cell where the sgRNA is detected. (right) Our strategy compares each sgRNA to all possible combinations of other sgRNAs targeting the same gene.

c.Schematic of our strategy to filter out poorly performing sgRNAs. Continuing from (b), we then calculate the energy distance between cells with a given sgRNA versus cells from all possible combinations of sgRNAs targeting the same gene. Next, we rank order these comparisons by energy distance. If a given sgRNA exhibits significant skewing in this rank ordering, then the sgRNA is an outlier.

d.Example outlier analysis for sgRNAs targeting HIF1A, where we successfully identify 62162262.23-P1P2-1 as an outlier.

**Extended Data Fig. 5**

a. Schematic of gene program analysis

b. Summary statistics for gene program analysis. (left) Shown is the distribution of the number of programs significantly associated with each TF perturbation. (right) Shown is the distribution of the number of TFs significantly associated with each gene program.

c. Feature plots showing: expression of an early CM marker (FN1), expression of a late CM marker (TNNT2), and GP1_car dev_ usage.

d. Volcano plot of TFs regulating GP1_car dev_. Known CHD TFs are highlighted in red.

e. Immunofluorescent staining of TNNT2 in iPSC-CM of RCOR2 KD, GATA4 KD.

f.Schematic of known CHAMP1 mutation in neuro developmental diseases and de novo mutations found in a CHD patient. Clinical information is from Jin et al^41^.

**Extended Data Fig. 6**

a. Example of a sub-network of CHD genes regulated by TFs. Red edges indicate up-regulation, and blue edges indicate down-regulation. Purple nodes indicate transcription factors, and green nodes indicate target CHD genes.

b. ChIP-Seq intensity on top 100 GP1_car dev_ promoter, and metaplot of the overall binding pattern.

c. Bubble plot showing the expression of the marker genes in non-targeting gRNA cells in time-course Perturb-Seq library. The size of the dot indicates fraction of cells expressing marker genes, and color indicates mean CPM in non-targeting cells.

d. Gene regulatory network showing transition of GP1_car dev_ regulating TFs at D4, D8, D12, D29 (top to bottom).

**Extended Data Fig. 7**

a. Heatmap showing top marker genes in each cardiac differentiation GP.

b. Bar plot showing hub TFs of each development-associated GP based on top in-degree nodes in TF-TF regulatory networks. Red color indicates activator of the GP whereas blue color indicates repressor of the GP.

c. Gene regulatory network of the Epicardial, cardiac mesoderm, FHF, SHF GPs (top to bottom)

**Extended Data Fig. 8**

a. Heatmap showing significant TF-GP regulation at various thresholds.

b. Heatmap showing significant OR gene (negative control)-GP regulation at various thresholds.

c. Heatmap showing the number of the maximum overlapping genes between cell-type annotation markers (CS8/CS9) and gene program top300 genes.

d. Schematic of in vivo differentiation lineage from epiblast.

e. Heatmap showing early development cell-type GP regulation by TF KD. Columns indicate CRISPRi targets and rows indicate GPs.

f. (left) Heatmap showing top marker gene in early development cell-type GPs. (right) Heatmap showing early development cell-type GP regulation by selected TF KD. Columns indicate CRISPRi targets and rows indicate GPs.

h. Barplot showing qPCR result of EOMES/TBXT (primitive streak/mesendoderm markers) upon ZBTB12/GTF2I/RCOR2 knock-down in Day 2 hESC-CMs.

i. Heatmap showing cardiac differentiation GP regulation by TF KD. Columns indicate CRISPRi targets and rows indicate GPs.

j. Schematic of in vivo differentiation lineage of cardiomyocyte subtypes.

k. (left) Heatmap showing top marker gene in cardiomyocytes subtype GPs. (right) Heatmap showing cardiomyocytes subtype GP regulation. Columns indicate CRISPRi targets and rows indicate GPs.

**Extended Data Fig. 9**

a. Schematic of the cardiac differentiation and marker genes(left), manually curated regulator(center), and their criteria (right)

b. Venn plot showing overlap between manually curated regulators and gene-program regulators on cardiac mesoderm regulation.

c. Venn plot showing overlap between manually curated regulators and gene-program regulators on cardiomyocytes regulation

**Extended Data Fig. 10**

a. Feature plots for TNNT2 (late cardiac marker) and FN1 (early cardiac marker).

b. Feature plot illustrating the cell distribution for each of the 26 Enhancer Perturb-Seq libraries.

c. Prioritization of target genes and enhancers based on energy distance statistics. Each dot represents a perturbation of promoters or enhancers. Perturbations with p<0.001 in energy statistics are shown in orange. 116 promoters and 194 enhancers are prioritized.

**Extended Data Fig. 11**

a. t-SNE embedding where each dot represents a perturbation in the Enhancer Perturb-Seq experiment. Colors indicate clusters. Enhancers are labeled as “Element” plus a numerical identifier (see **Supplementary Table 19**).

b. (top) Zoomed-in view of HOXB4 promoters and enhancers cluster in the embedding and genomic positions of these elements. ATAC-Seq is from Day12 WTC11-derived cardiomyocytes and H3K27ac is from D15 H9-derived cardiomyocytes^20^. (bottom) qPCR validation of downstream gene expression changes after promoter/enhancer CRISPRi.

c. (top) Zoomed-in view of TBX5 promoters and enhancers cluster in the embedding and genomic positions of these elements. (bottom) qPCR validation of downstream gene expression changes after promoter/enhancer CRISPRi.

d. Schematic of normalized energy distance. In this example, if the energy distance between background and promoter perturbation is 20 and the energy distance between promoter and enhancer perturbation is 8, then the normalized energy distance for the enhancer perturbation is 0.4.

e. Distribution of normalized e-dist between TF-OR genes and TF-neighbor regulatory elements. (OR: olfactory receptor genes, which serve as negative controls)

f. The proportion of identified enhancers in different metrics by target-enhancer distance. A number of enhancer hits are shown at the top of each bar.

g. Impact of all perturbed enhancers on normalized target gene expression and transcriptome-wide energy distance. Color indicates the hits of enhancers found by different metrics.

h. Venn diagram of enhancers identified by differential expression analysis (DEG, green), compared to our transcriptome-wide approach (blue).

**Extended Data Fig. 12**

a. Schematic architecture of Percoder.

b. Bar plot showing Percoder’s prediction accuracy depends on the number of cells used in the input cell sets.

c. Example ROC curves of key cardiac TFs, GATA4.

d. Venn diagram showing shared perturbations from two batches of Perturb-Seq experiments: TF Perturb-Seq (Fig. 1) and enhancer Perturb-Seq (Fig. 5).

e. Schematic of out-of-sample prediction. External datasets (other Perturb-Seq experiments or patient-derived samples) are integrated into the shared embeddings with the TF Perturb-Seq dataset. We then use the Percoder classifier to predict dysregulated TFs in the set of cells from the external single-cell RNA-Seq dataset.

f. Top k prediction accuracy for shared perturbations between two Perturb-Seq batches. Blue bar indicates accuracy in the classifier and orange bar is the result from a random classifier.

g. Percoder prediction accuracy using the out of sample Enhancer Perturb-Seq dataset, when using PCA (blue) or scVI (orange) embeddings.

h. Top k prediction accuracy for shared perturbations between two Perturb-Seq batches using Geneformer embedding and cosine similarity.

i. Histogram showing the rank of actual perturbations among the top differentially regulated regulons identified by SCENIC analysis in perturbed cells of training dataset.

j. Histogram showing the effects on corresponding regulons by actual perturbations in perturbed cells of training dataset.

k. We applied Percoder on scRNA-Seq data derived from hypoplastic right heart syndrome patient-derived iPSCs differentiated to cardiomyocytes^89^. The volcano plot shows Percoder predictions of dysregulated regulators. Color indicates the Percoder score in the patient sample (not normalized relative to control).

**Extended Data Fig. 13**

Feature plots illustrating the quality of single-cell data integration with (left) PCA and (right) scVI-based UMAP. Shown are the cells from two Perturb-Seq experiments (Figure 1 data; Figure 5 data), patient-derived cardiomyocytes from three published studies (Xu, Gonzalez, Yang), and two marker genes (TNNT2, FN1).

## SUPPLEMENTARY TABLES

**Supplementary Table 1:** List of sgRNAs used in TF Perturb-Seq screen.

**Supplementary Table 2:** Key statistics of TF Perturb-Seq libraries.

**Supplementary Table 3:** Bulk sgRNA count at D0, D7, and D12 (from Perturb-Seq) of differentiation.

**Supplementary Table 4:** Outlier analysis of sgRNAs in TF Perturb-Seq screen.

**Supplementary Table 5:** Filtered list of non-targeting gRNAs used for background

**Supplementary Table 6:** Permutation analysis for sgRNAs in TF Perturb-Seq screen.

**Supplementary Table 7:** Overlap of energy distance hits with de novo CHD variants.

**Supplementary Table 8:** Summary of energy distance TF hits.

**Supplementary Table 9:** Top 300 genes in each gene program identified by cNMF.

**Supplementary Table 10:** Gene programs associated with at least 1 TF perturbation.

**Supplementary Table 11:** Gene Ontology terms associated with each gene program.

**Supplementary Table 12:** List of sgRNAs used in Time-course TF Perturb-Seq screen.

**Supplementary Table 13:** Top 300 genes in each gene program at D4, D8, D12, D29 differentiation identified by cNMF.

**Supplementary Table 14:** Gene programs associated with at least 1 TF perturbation at D4 Time-course TF-Perturb-Seq.

**Supplementary Table 15:** Gene programs associated with at least 1 TF perturbation at D8 Time-course TF-Perturb-Seq.

**Supplementary Table 16:** Gene programs associated with at least 1 TF perturbation at D12, Time-course TF-Perturb-Seq.

**Supplementary Table 17:** Gene programs associated with at least 1 TF perturbation at D29 Time-course TF-Perturb-Seq.

**Supplementary Table 18:** Summary of TF-TF interaction analysis.

**Supplementary Table 19:** List of sgRNAs used in Enhancer Perturb-Seq screen.

**Supplementary Table 20:** Key statistics of Enhancer Perturb-Seq libraries.

**Supplementary Table 21:** Filtered list of non-targeting gRNAs used for background.

**Supplementary Table 22:** Outlier analysis of sgRNAs in Enhancer Perturb-Seq screen.

**Supplementary Table 23:** Permutation analysis for sgRNAs in Enhancer Perturb-Seq screen.

**Supplementary Table 24:** Summary of enhancer hits.

**Supplementary Table 25:** Summary of Promoter-Enhancer relative e-dist analysis.

**Supplementary Table 26:** Summary of Geneformer prediction statistics.

**Supplementary Table 27:** Summary of SCENIC regulon changes by actual perturbed cells.

**Supplementary Table 28:** Summary of Percoder predictions in Xu et al dataset.^88^

**Supplementary Table 29:** Summary of Percoder predictions in Gonzalez-Teran et al dataset.^60^

**Supplementary Table 30:** Summary of Percoder predictions in Yu et al dataset.^90^

**Supplementary Table 31:** Primers used in this study.

