## Extended Figures for "Regulatory network hubs guide dynamic human lineage specification"

#### Extended Data Fig. 1

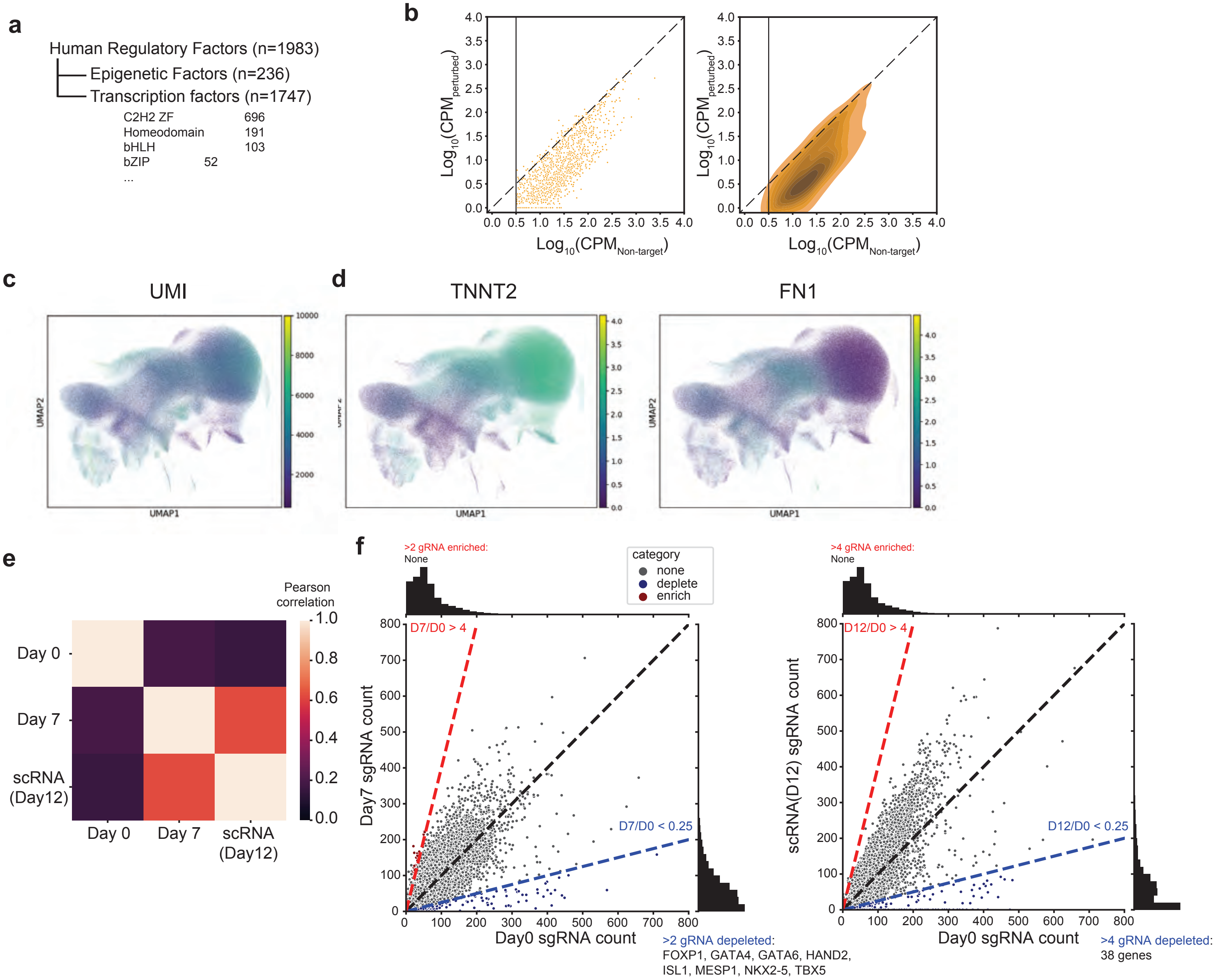

### Extended Data Fig. 2

a

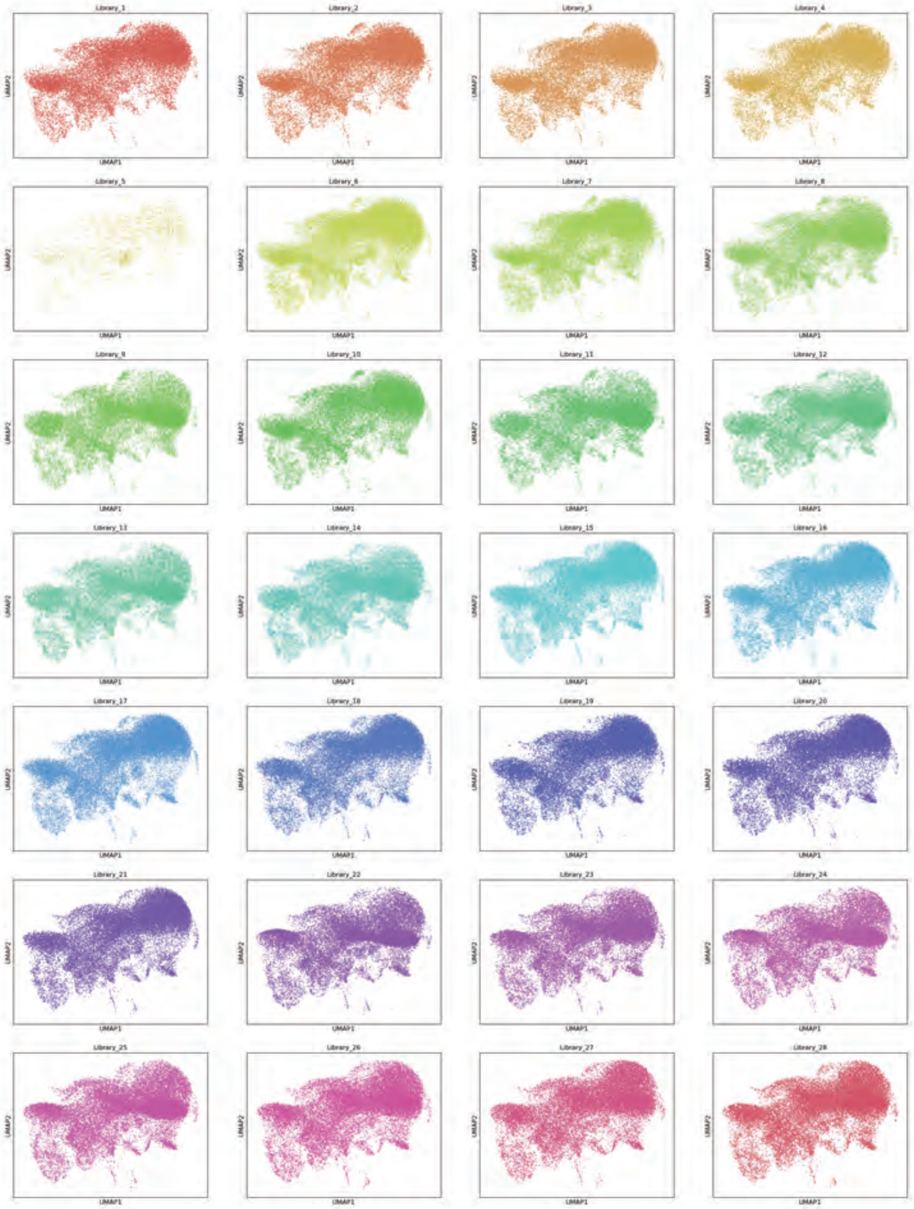

#### Extended Data Fig. 3

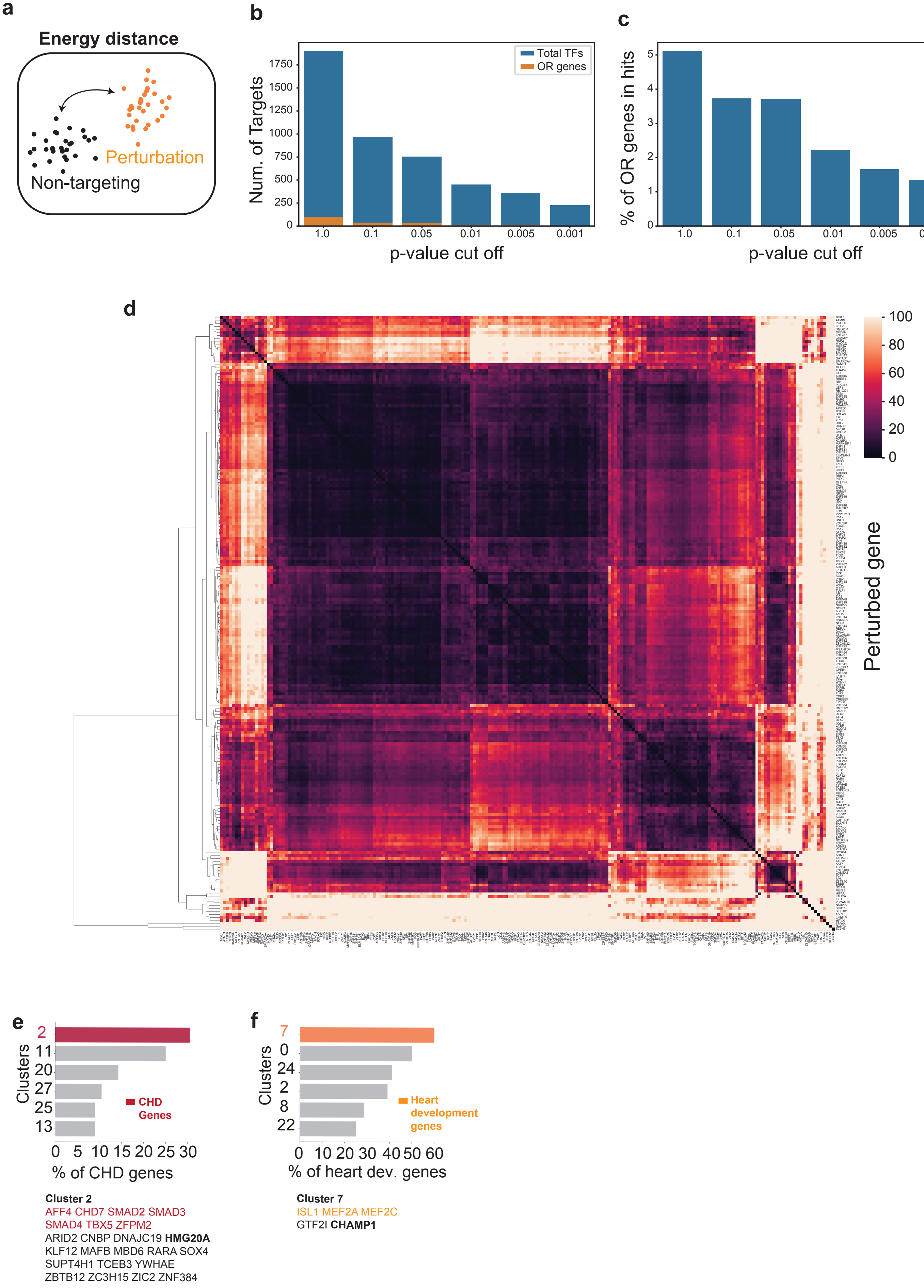

**Extended Data Fig. 4**

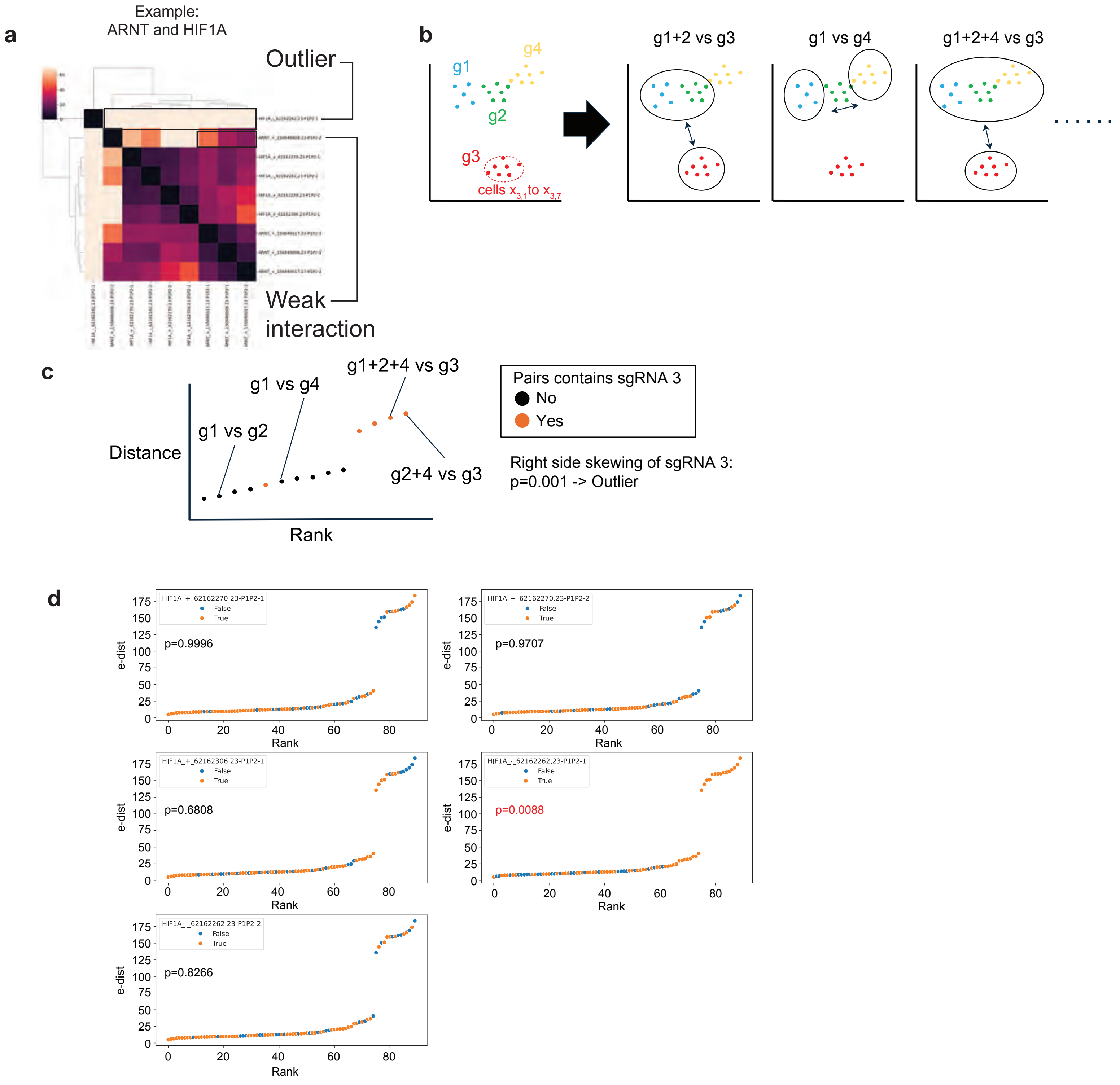

#### Extended Data Fig. 5

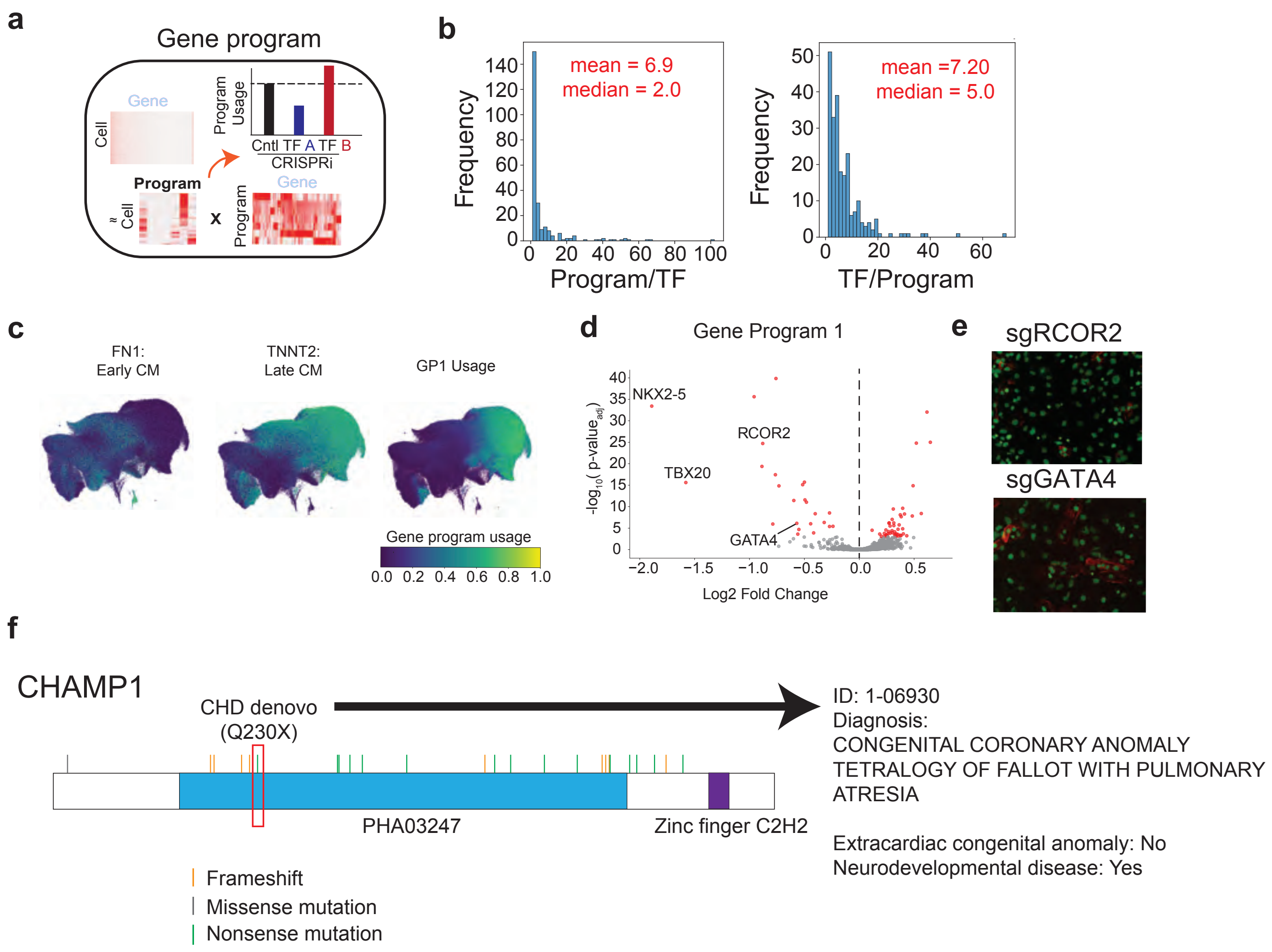

#### Extended Data Fig. 6

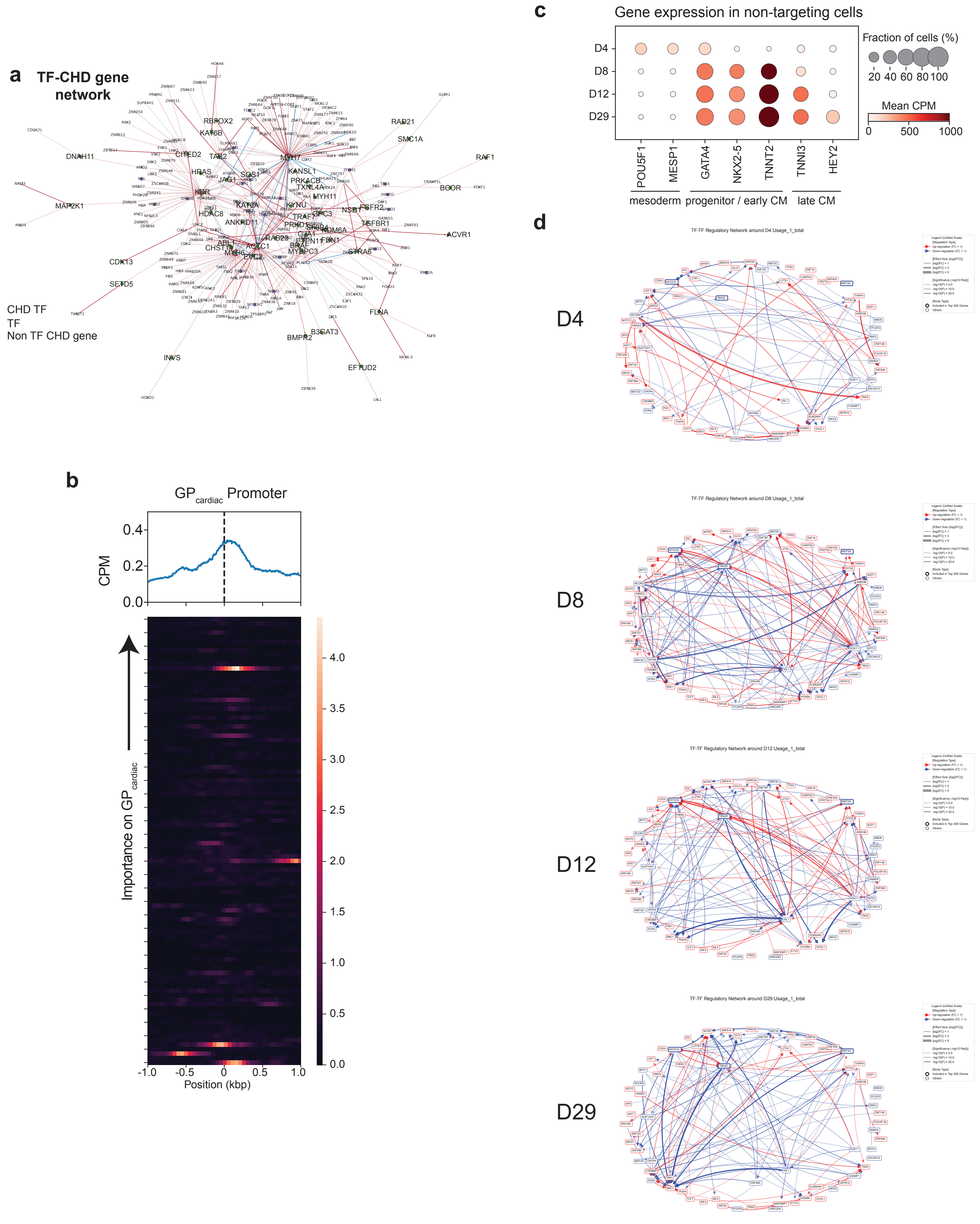

### Extended Data Fig. 7

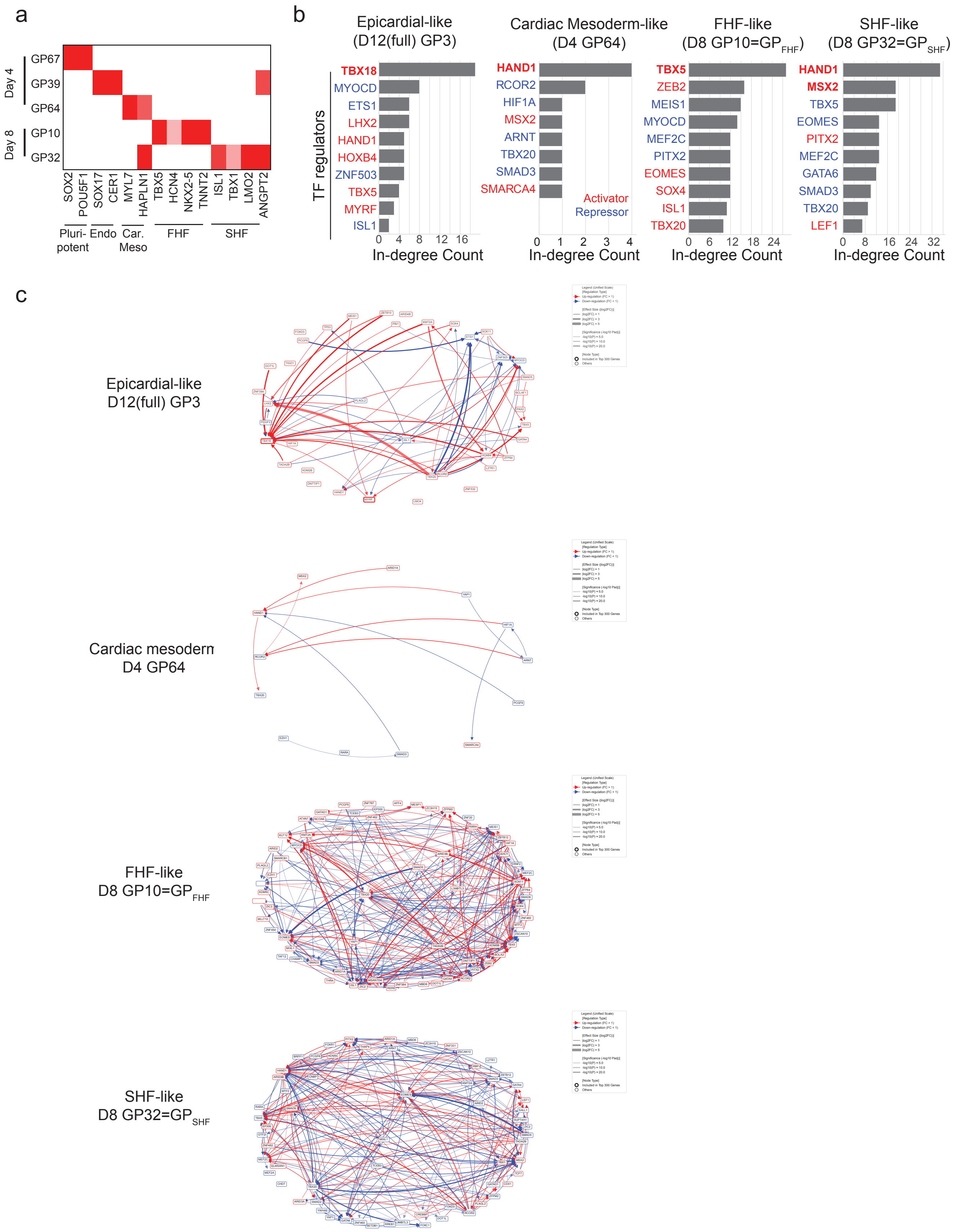

### Extended Data Fig. 8

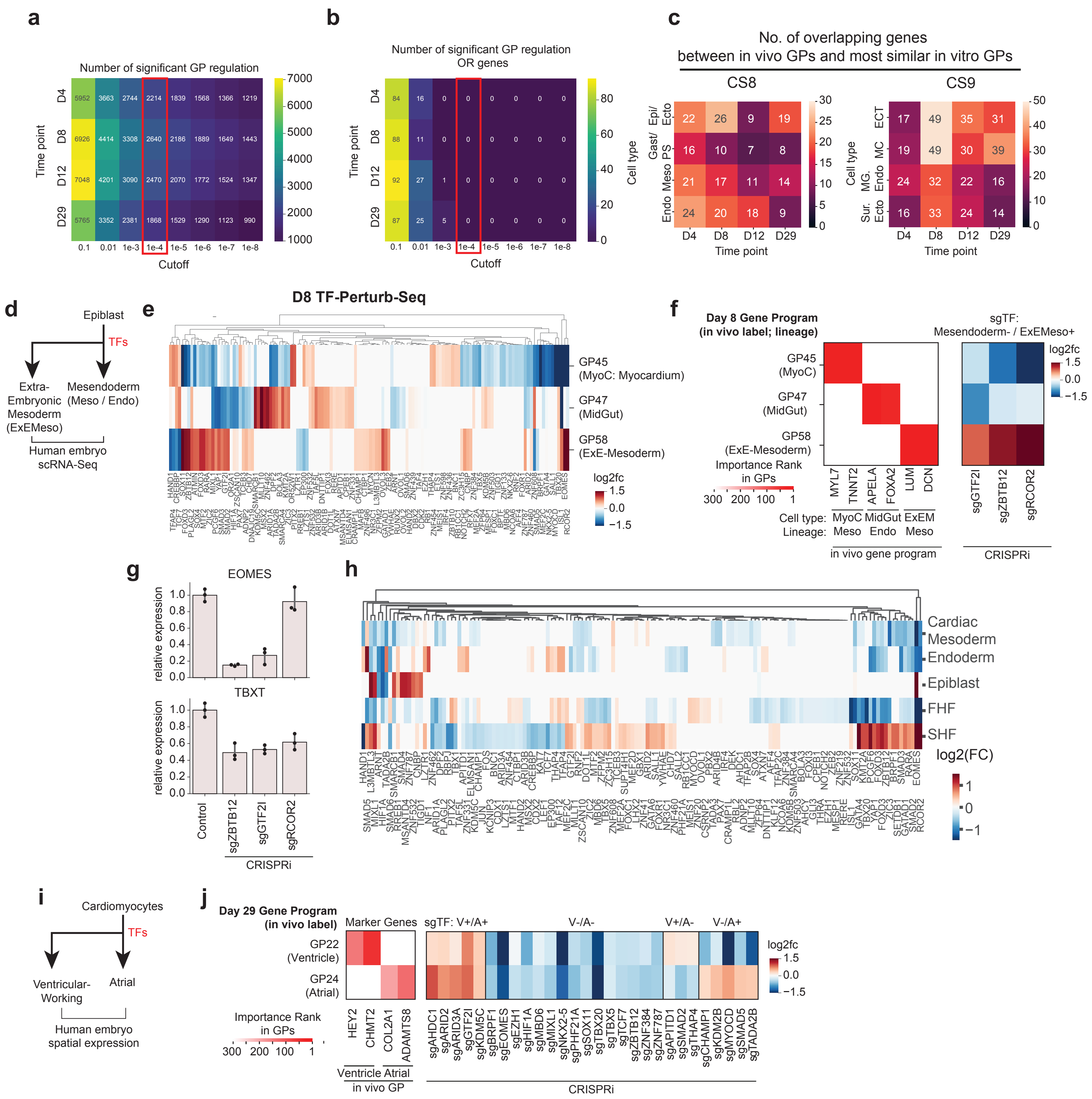

#### Extended Data Fig. 9

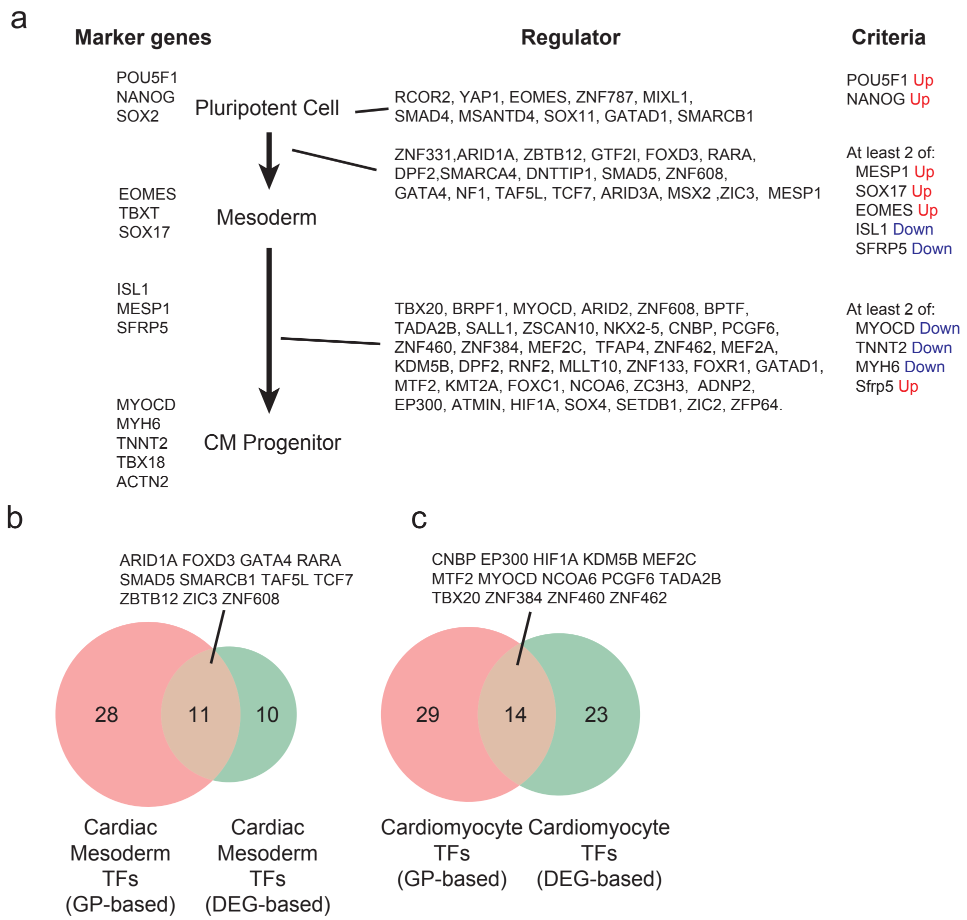

**Extended Data Fig. 10**

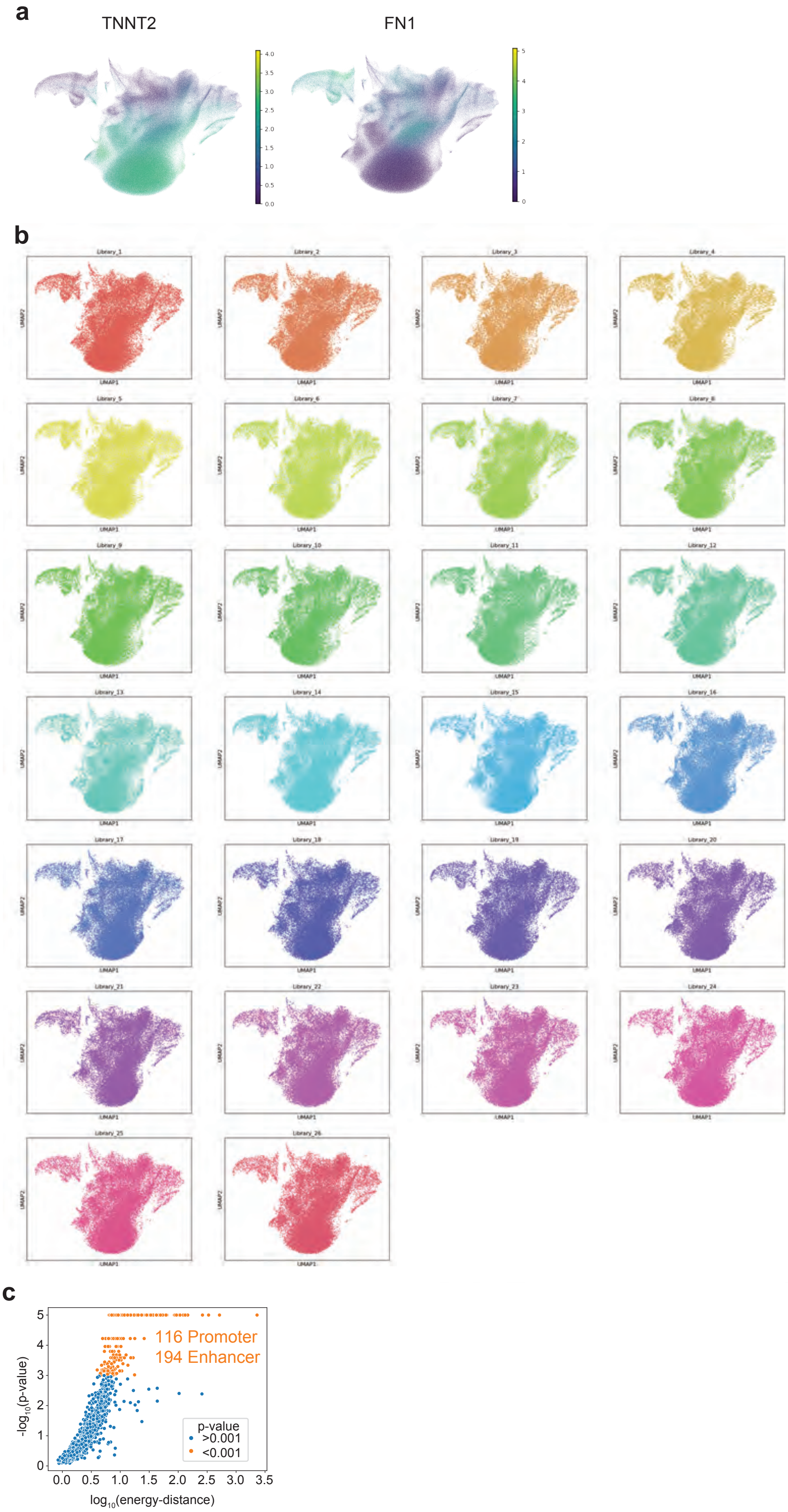

### Extended Data Fig. 11

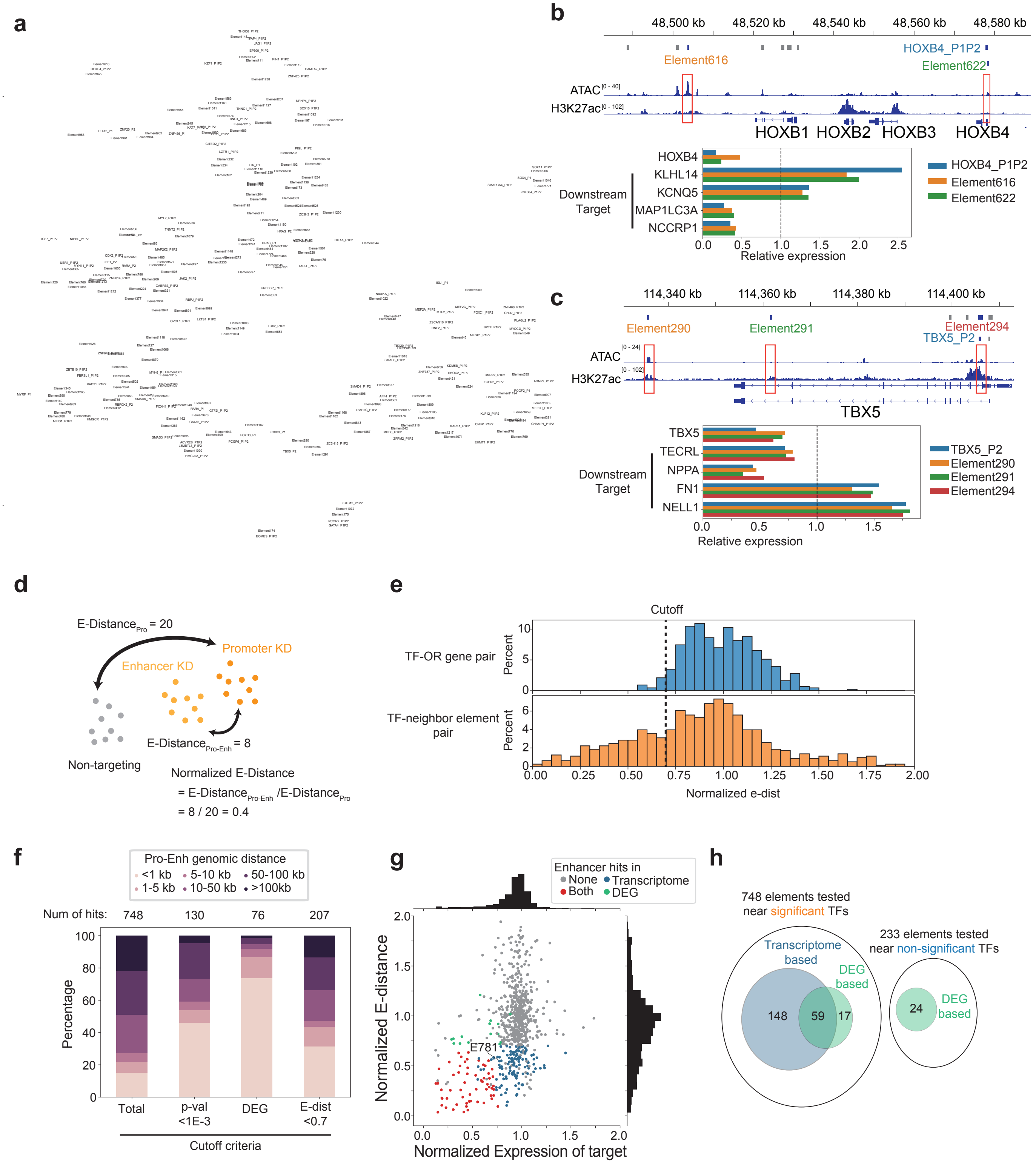

#### Extended Data Fig. 12

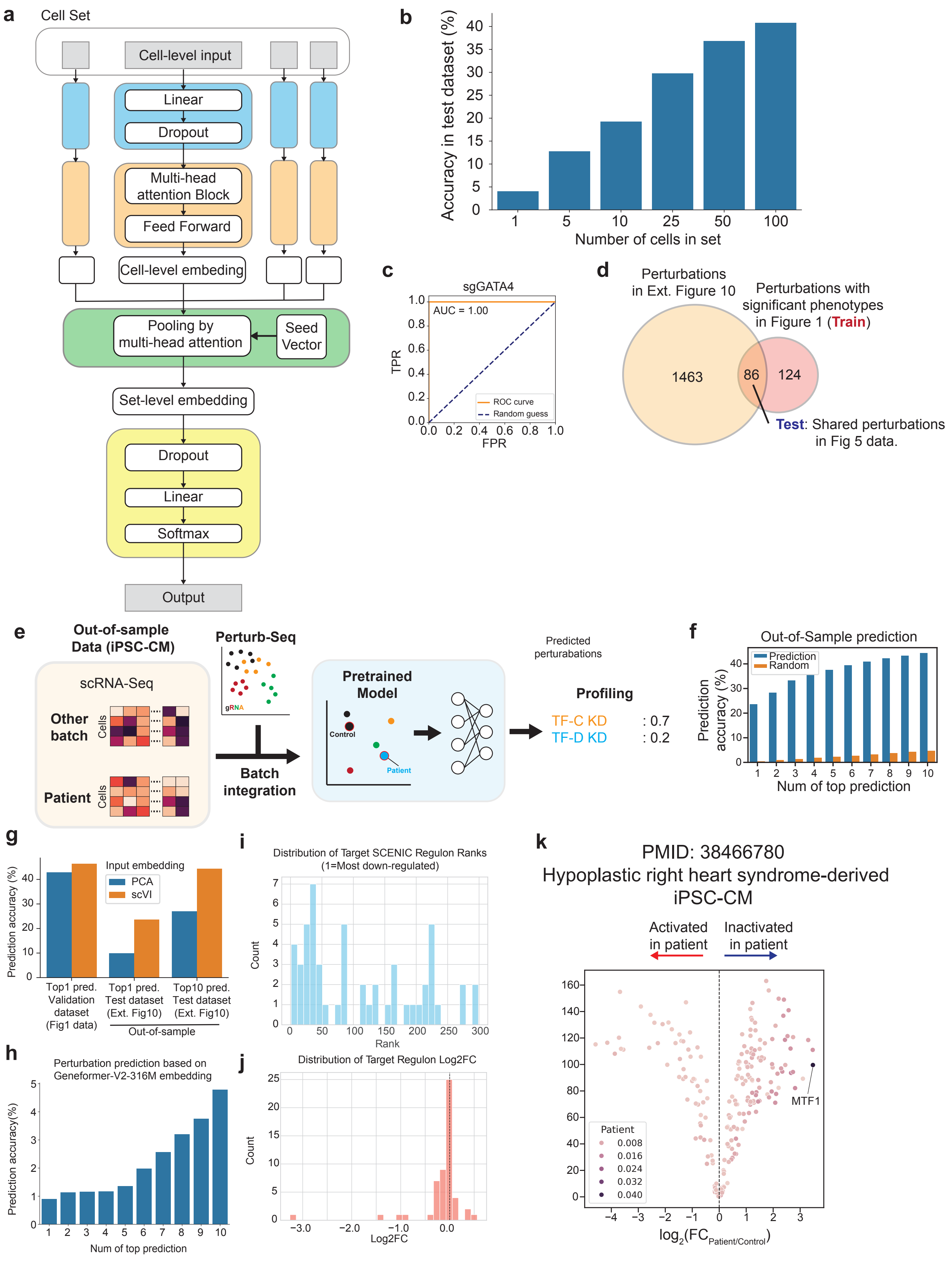

#### Extended Data Fig. 13

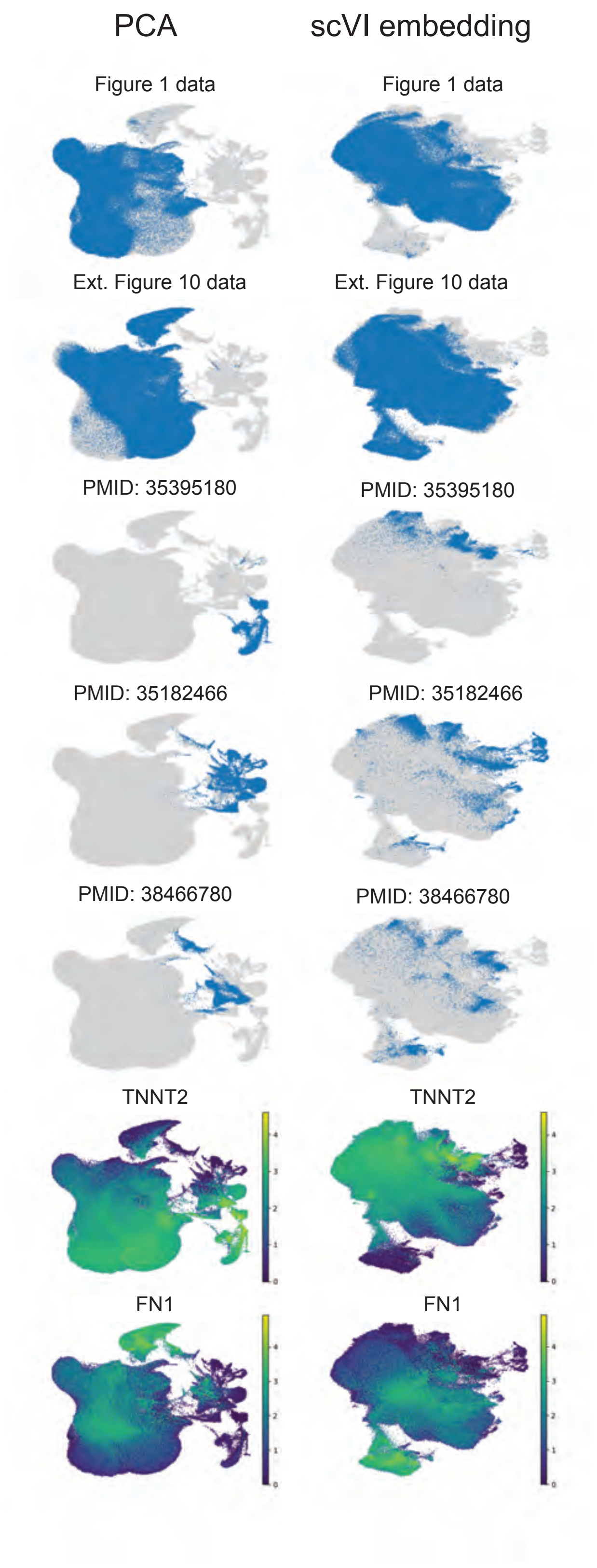
